# Temporal coordination of tissue transformation, olfactory sensory neural development and central axon projections through morphogens

**DOI:** 10.1101/2025.07.21.666031

**Authors:** Shao-Chieh Chen, Tsai-Ming Lu, Chun-Ting Lin, Iris Low, Ya-Hui Chou

**Author notes:** **Conflict of interest** The authors declare no competing interests.

## Abstract

During development, sensory neurons arise in the peripheral sensory organ in close spatial and temporal coordination with the dynamic morphological transformation of the sensory organ in the central nervous system. Moreover, intricate coordination exists between the peripheral map and the central map. Yet, it remains largely unclear what mechanisms are responsible for orchestrating such coordination and whether these mechanisms might be evolutionarily conserved. Here, we performed a systematic analysis of the sensory organ transformation process, defining the expression patterns of morphogens and their receptors at multiple developmental stages and creating receptor mutants in sensory neurons and projections. These experiments revealed combinatorial codes of morphogens that are utilized to coordinate sensory circuit development. Remarkably, two distinct strategies were likely deployed by different morphogen families, including a two-step strategy (same ligand from different sources at two different stages) and a ligand switch strategy (different ligands at two different stages).

**Significance Statement:** Tissue transformation, dissociation and integration occur in all species. However, it remains largely unclear how such transformations are coordinated with cell fate determination, local cell migration and axonal guidance both temporally and spatially. We found *Drosophila* antennal discs undergo a series of tissue transformation steps to develop a three-dimensional structure from a two-dimensional neuroepithelium. During this tissue transformation, olfactory sensory neurons acquire their distinct cell types and project their axons to specific targets in the antennal lobe. The whole process is temporally and spatially coordinated by combinatorial codes of morphogens.

## Introduction

Sensory neurons arise in the peripheral sensory organ in tight coordination with the dynamic morphological transformation of the sensory organ during development. This intricate developmental process often involves a nearly two-dimensional epithelium or primordium transforming into a three-dimensional sensory organ, while sensory neurons simultaneously become identifiable in peripheral tissues and extend axons to the central nervous system. The process requires precise temporal and spatial coordination, and it exhibits further complexity due to tissue segregation, migration and integration into one particular sensory organ. However, the mechanisms responsible for orchestrating such coordination remain largely undefined, and it is also unclear whether these mechanisms might be evolutionarily conserved.

In the olfactory circuit, olfactory sensory neurons (OSNs or ORNs) originate from peripheral tissue (i.e., the olfactory epithelium in rodents and antennae/maxillary palps in insects). The axons of OSNs/ORNs expressing the same odorant receptors converge into bundles and project to specific glomeruli in the primary olfactory information processing center in the brain (i.e., the olfactory bulb in rodents and antennal lobe in insects) (Barnum and Hong 2022). Interestingly, the somas of different classes of olfactory sensory neurons in the olfactory epithelium or antennae exhibit a coarsely defined topographical distribution (Miyamichi et al. 2005; Shanbhag et al. 1999), which is coordinated to a certain degree with their central axon targeting sites (Couto et al. 2005; Zhu et al. 2022). Hereafter, the topographic distribution of OSN/ORN somas in the peripheral sensory organ will be referred to as the “peripheral map,” while the axon projections in the olfactory bulb or the antennal lobe will be called the “central map.”

Rodent odorant receptors mediate activity-dependent axon targeting to specific glomeruli of the olfactory bulb (Imai et al. 2006; Takeuchi and Sakano 2014; Serizawa et al. 2006). However, *Drosophila* odorant receptors are not involved in axon targeting (Dobritsa et al. 2003; Larsson et al. 2004). *Drosophila* Notch coordinates the peripheral map and central map by defining cell fates of sensory neurons and controlling specific axon guidance molecules (Joo et al. 2013; Endo et al. 2007). Moreover, Hedgehog (Hh) contributes to the positioning of sensory neuron somas within peripheral sensory organs and underlies the establishment of a coordinated sensory map (Chou et al. 2010). Nevertheless, the somas of a given class of ORNs can be scattered within a certain range after sensory organ transformation (Song et al. 2012), and the relative positioning of a given class of ORN somas in the periphery is positively but weakly correlated with the positioning of central axon-targeted sites (Chai et al. 2019).

While extensive research has been conducted on the development of ORNs and their axon targeting (Sweeney et al. 2007; Sweeney et al. 2011; Hong et al. 2009; Ward et al. 2015; Hummel et al. 2003; Kaur et al. 2019; Sakurai et al. 2009; Endo et al. 2011; Ray et al. 2007; Ang et al. 2003; Silbering et al. 2011; Fishilevich and Vosshall 2005; Couto et al. 2005; Barish et al. 2018; Rodrigues and Hummel 2008; Lattemann et al. 2007), a critical knowledge gap exists regarding the temporal coordination between development of peripheral sensory organs from two-dimensional to three-dimensional structures and the development of the peripheral sensory neuron soma map. Morphogens regulate differentiation and neural identities in neural plate (Sagner and Briscoe 2017), patterns fly eye and wing in development (Sagner and Briscoe 2017; Kumar 2001; Rossi et al. 2021; Vincent and Briscoe 2001), and involve in mouse and C. elegans axon targeting (Liao et al. 2018; Klassen and Shen 2007; Charron et al. 2003). Yet, systematic explorations of their involvement in neural map formation and coordination are lacking. Here we utilize *Drosophila* as a model system to investigate whether and how morphogens may function to coordinate three critical developmental aspects: (1) transformation of a sensory organ from a two-dimensional to three-dimensional structure, (2) the soma positions of sensory neurons, and (3) axon targeting of sensory neurons in the central brain.

## Materials and Methods Materials

### Drosophila melanogaster

Flies were raised in corn meal at 25°C in an incubator with 12 h-12 h light-dark control. Both males and females were used for experiments with roughly equal numbers. Flies used in this paper were *Canton-S* (Bloomington *Drosophila* Stock Center, BDSC 1), *p{PZ}hh[P30]* (BDSC 5530), *UAS-mCD8GFP.1/CyO* (Lee and Luo 1999), *p{GAL4-dpp.blk1}c40.1* (Gift from Dr. Henry Sun), *UAS-GFP.nls (14)* (BDSC 4775), *ptc-GAL4.559.1*(BDSC 2017), *NP2211* (Kyoto Stock Center 112825), *bnl[MI00874-GFSTF.1]* (BDSC 61653), *Actbeta[MI14795-DH.PT-GFSTF.1]* (BDSC 93676), *NP2048* (Kyoto Stock Center 104070), *vn[MI05869-GFSTF.1]* (BDSC 60234), *P{ PZ}vn[10567]* (BDSC 11749), *NP6034* (Kyoto Stock Center 113819), *w[*] peb-GAL4; FRTG13 UAS-mCD8GFP* (gift from Dr. Liqun Luo), *AM29-GAL4* (Gift from Dr. Chihiro Hama), *Gr21a-GAL4 (D1)* (Gift from D. Kristin Scott), *P{Gr21a-GAL4.9.323}* (BDSC 57600), *Or10a-GAL4 (14.1)* (Gift from Dr. Liqun Luo), *Or10a-GAL4 (15)* (Gift from Dr. Liqun Luo), *Or22a-GAL4 (14.2)* (BDSC 9951), *Or22a-GAL4 (III)* (Gift from Dr. Larry Zipursky), *P{Or23a-GAL4.7.818}17.2* (BDSC 9955), *P{Or23a-GAL4.7.818}17.4* (BDSC 9956), *P{Or42a-GAL4.F}48.3B* (BDSC 9970), *Or42a-GAL4 (43.4)* (Gift from Dr. Liqun Luo), *Or43a-GAL4 (18d)* (Gift from Dr. John Carlson), *Or46a-GAL4 (III)* (Gift from Dr. Larry Zipursky), *Or47a-GAL4 (15.2)* (Gift from Dr. Leslie Vosshall), *P{Or47a-GAL4.8.239}15.4A* (BDSC 9982), *P{Or47b-GAL4.7.467}15.6* (BDSC 9984), *P{Or47b-GAL4.7.467} Bacc15.5A* (BDSC 9983), *Or59c-GAL4 (21.7)* (Gift from Dr. Leslie Vosshall), *P{Or59c-GAL4.C}129t5.4, w\** (BDSC 23900), *Or67b-GAL4 (B3960)* (Gift from Dr. John Carlson), *Or67b-GAL4 (B3960.1)* (Gift from Dr. John Carlson), *Or71a-GAL4 (III)* (Gift from Dr. Liqun Luo), *Or71a-GAL4 (30.3)* (BDSC 23121), *Or83c-GAL4 (1)* (Gift from Dr. John Carlson), *Or85e-GAL4 (1.14.1)* (Gift from Dr. John Carlson), *P{Or85e-GAL4.W}2.19.1* (BDSC 23291), *Or88a-GAL4 (X2db)* (Gift from Dr. John Carlson), *Or92a-GAL4 (38.1)* (Gift from Dr. Liqun Luo), *P{Or92a-GAL4.F}62.4* (BDSC 23140), *smo[3]* (Chou et al. 2010), *ptc[IIW]*(Chou et al. 2010), *tkv[strII]* (Gift from Dr. Henry Sun), *fz[H51]* (Gift from Dr. Makoto Sato), *fz2[C1]* (Gift from Dr. Makoto Sato), *fz[H51] fz2[C1]* (Gift from Dr. Makoto Sato), *FRT42D EGFR[-]* (Gift from Dr. Henry Sun), *W[*]; FRTG13 babo[Fd4] GH146-GAL4/CyO* (Gift from Dr. Liqun Luo), *wit[G5] FRT2A* (Gift from Dr. Liqun Luo), *dSMAD2[-] UAS-mCD8GFP FRT19A/FM7c* (Gift from Dr. Liqun Luo), *W[*]; drl[343] FRT40A/CyO* (Gift from Dr. Liqun Luo), *hsFLP122; FRTG13 GH146-GAL4 UAS-mCD8GFP/CyO* (Gift from Dr. Liqun Luo), *y w hsFLP UAS-mCD8GFP; tubp-GAL80 FRT40A GH146-GAL4/CyO*(Chou et al. 2010), *y w hsFLP UAS-mCD8GFP; FRT42D ptc[IIW] GH146-GAL4 UAS-mCD8GFP/CyO* (Chou et al. 2010), *w [*]; tubGAL80 FRT40A (Lee and Luo 1999), w[*]; FRT42D tubp-GAL80/CyO* (Lee and Luo 1999), *w[*];tubp-GAL80 FRT2A/TM6B Tb* (Lee and Luo 1999), *GH146-GAL4* (Gift from Dr. Liqun Luo). Detailed genotypes were listed in Table S1.

### Antibodies

The primary antibodies used in this study were rabbit anti-β-Galactosidase (08559762, MP Biomedicals), mouse anti-Wg (4D4, Developmental Studies Hybridoma Bank (DSHB)), rat anti-DN-cadherin (DN-Ex#8, DSHB), mouse anti-Bruchpilot (nc82, DSHB), chicken anti-GFP (GFP-1020, Aves Labs), rabbit anti-GFP (A11222, Invitrogen), rat anti-Spitz (Anti-Spitz, DSHB), rat anti-mCD8 (MCD0800, Invitrogen). The secondary antibodies used and from Jackson ImmunoResearch Laboratories were goat anti-mouse IgG-Alexa Fluor® 488 (115-545-166), goat anti-mouse IgG-CyTM3 (115-165-166), goat anti-mouse IgG-Alexa Fluor® 647 (115-605-166), goat anti-rabbit IgG-Alexa Flour® 488 (111-545-144), goat anti-rabbit IgG-Cy3 (111-165-144), goat anti rabbit IgG-Alexa Fluor® 647 (11-605-144), goat anti-rat IgG-Alexa Fluor® 488 (112-545-167), goat anti-rat IgG-CyTM3 (112-165-167), goat anti-rat IgG-Alexa Fluor® 647 (112-605-167), goat anti-rat IgG-DyLight 649 (112-495-167), and goat anti-chicken IgY (IgG)-DyLight488 (103-485-155).

### Experimental design and statistical analyses

#### Dissecting early pupal discs

To maintain disc orientation along the body axes, the anterior cuticle of early pupae was carefully cut by surgery scissors along around 1/3 of the body in 4% paraformaldehye, followed by a cut from the ventroposterior to the mouth hook. The fat body and debris were gently removed by washing the cuticle in fixative solution. The cuticle with attached imaginal discs was subjected to immunostaining (see below).

#### Immunohistochemistry

Brains, eye-antennal discs or adult heads (carrying antennae and maxillary palps) of larvae, pupae or adults were dissected in freshly prepared cold 4% paraformaldehyde/ PBS and fixed in the same fixative solution on an ice bath immediately after dissection. This step was followed by a fixation at room temperature (RT) for 20 min. Samples were washed with PBST (0.3% Triton X in PBS) at RT for 20 min three times to remove fixative and then incubated in blocking solution [5% normal goat serum (NGS) in PBST] at RT for 30 min. Samples were then incubated with primary antibodies in 5% NGS/PBST at 4°C for 2-3 days. Samples were washed with PBST three times as described above and then incubated with diluted secondary antibodies (1:500 in PBST) at 4°C overnight. Next, the tissues were washed with PBST three times as described above and then incubated with mounting solution (SlowFade^TM^ Gold Antifade Mountant, S36936, Invitrogen) at 4°C at least overnight before being mounted to the slide.

The dilutions of primary antibodies were as follows: rabbit anti-b-GAL (1:1000), mouse anti-Wg (1:50), mouse anti-Brp (1:25), rat anti-mCD8 (1:100), Chicken anti-GFP (1:500), rabbit anti-GFP (1:1000), rat anti-Spitz (1:40), goat anti-Activin (1:40). The dilutions of all secondary antibodies were 1:500.

#### Confocal imaging and live imaging

Samples were imaged using a Zeiss LSM 510, a Zeiss LSM780 equipped with Mai-Tai HP-1040 (Spectra-Physics), or a Zeiss LSM700 confocal microscope. Images were processed using AimImageExaminer (Carl Zeiss), Zen black (Carl Zeiss), Fiji [https://fiji.sc], and Adobe Photoshop (Adobe). Live images of antennae were acquired with a Zeiss Axiovision 8 microscope, equipped with Pentagon FC Camera (Pentagon), and processed using PTImage analsis software (Pentagon).

#### Statistical analyses

No samples were excluded, except for brains with obvious damage introduced during sample preparation. For eyFLP-induced *smo* or *smo tkv* MARCM clones, less than 10% of adult flies with obviously smaller antennae and/or maxillary palps were excluded from the dissection (Figure 6, Figure S4).

#### Code/Software

The software used in this study were Zen Black (Carl Zeiss), AimImageExaminer (Carl Zeiss), Fiji (https://fiji.sc), PTImage Analysis Software (Pentagon), R package Seurat (Version 5.1.0, CRAN), R package ggplot2 (Version 3.5.1, CRAN), Illustrator (Adobe), Photoshop (Adobe), BioRender (BioRender).

### Reanalysis of published data

#### Data processing and visualization of single-cell RNA-seq

The annotated single-cell RNA-seq datasets for the olfactory receptor neuron (ORN) and projection neuron (PN) classes were retrieved from the NCBI Gene Expression Omnibus (accession numbers: GSE162121, GSE161228). The preprocessing steps followed those described in previous studies (McLaughlin et al. 2021; Xie et al. 2021). The read count normalization method was as described in the aforementioned studies. The coordinates for t-Distributed Stochastic Neighbor Embedding (t-SNE), the cell type annotations, normalized expression levels, and developmental stage information were directly adopted from the source datasets. Using this approach, the figures presented in previous studies could be reproduced. The t-SNE plots were constructed using the DimPlot function in the R package Seurat (Hao et al. 2024), while the FeaturePlot function was used to generate feature plots showing expression profiles of individual genes. Dot plots were generated using the R package ggplot2. For dot plots, the average expression levels of specific genes were computed for each cell type as previously annotated. A gradient of expression levels from low to high is displayed in the figures instead of the exact values. The “Percent Expressed” metric indicates the proportion of cells within a given cell type that express a specific gene.

## Results

### *Drosophila* olfactory sensory organs undergo tissue dissociation, transformation, migration and integration during metamorphosis

*Drosophila* olfactory sensory organs include antennae and maxillary palps. The sac-like structures of antennae and maxillary palps develop from an epithelial antennal disc through the pupal stage (Postlethwait and Schneiderman 1971; Cohen and Jurgens 1989). The antennal disc is a part of eye-antennal disc, which is attached anteriorly to the larval mouth hook and posteriorly to the optic lobe (Figure 1A, B, left). As such, the antennal disc must dissociate from the eye disc, followed by further dissociation of the future antennal tissue and maxillary palp tissue. During this process, the antennal disc also migrates toward the developing labial disc so that maxillary palp can attach to the proboscis (Figure 1A, B, middle and right). In addition, the antennal disc of late 3rd instar larvae has been proposed to undergo 90° rotation during its morphological transformation at the pupal stage to establish sac-like antennae in adults (Postlethwait and Schneiderman 1971). How these developmental processes are coordinated spatially and temporally during pupal stage remains unclear.

**Figure 1.**
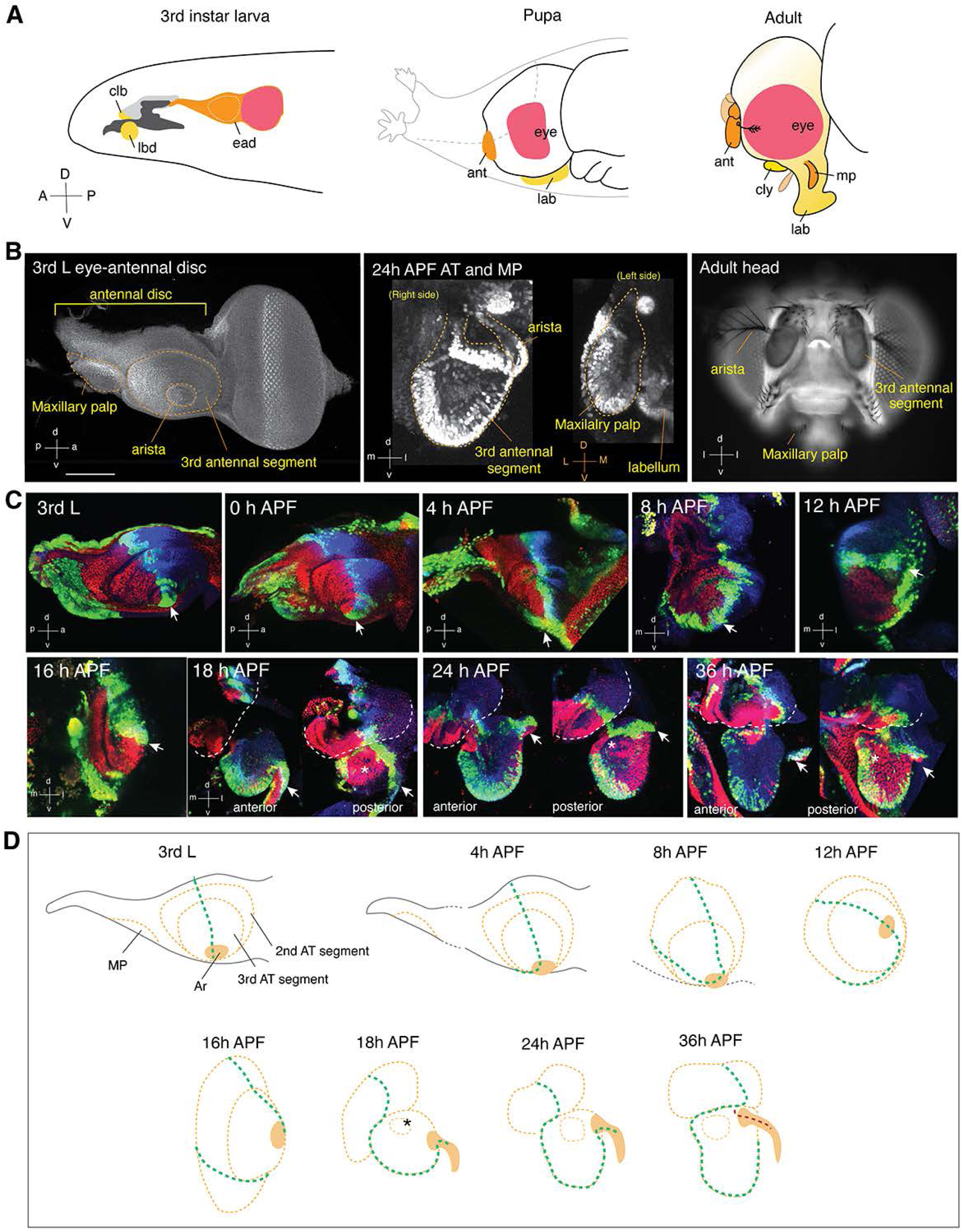
Morphological transformation of the olfactory sensory organ. (A) Schematic diagram of the relative positions of discs and appendages. (Left) Third instar larva. (Middle) Pupa. (Right) Adult. All images are aligned with same orientation: anterior left and dorsal up in lateral view. The eye-antennal disc contributes to the upper part of the proboscis, maxillary palps, antennae, head capsule and eyes. The labial disc contributes to the distal part of proboscis, including labellum. The clypeolabral disc (bud) becomes clypeolabrum in adult. ead: eye-antennal disc; clb; clypeolabral disc (labrum); cps, cephalopharyngeal skeleton (dark grey); dp, dorsal pouch (light grey); lbd, labial disc; lab, labellum; ant, antenna; mp, maxillary palp; cly, clypeus. (B) Confocal images (left and middle) and photograph (right) of larval eye-antennal disc, pupal antenna and maxillary palp, and adult head, respectively. Scale bar, 100 μm. A/a, anterior; P/p; posterior; D/d, dorsal; V/v, ventral; m, medial. To better illustrate the coordination of global and regional body plans, the body axes are designated with capital letters and disc/appendage axes are in lowercase. (C) Confocal images of antennal disc and antenna at different developmental stages. The posterior, midline (anterior-posterior boundary) and dorsal compartments of the antennal discs/antennae were labeled by *hh-lacZ* (red), *ptc-GAL4* driven nuclei-GFP (green), and Wg (blue), respectively. Arista and sacculus are indicated by arrows and asterisks, respectively. (D) Schematic diagram of antennal disc transformation. See text for details.

To address this question, we first examined the morphological changes of larval antennal discs during pupal stages (Figure 1C). Morphogens are known to be expressed in different compartments of larval antennal discs (Chou et al. 2010; Lebreton et al. 2008; Royet and Finkelstein 1997); for instance, Hedgehog (Hh) and patched (Ptc) are respectively expressed in the posterior compartment and at the boundary of anterior and posterior compartments (midline) of the antennal disc, while Wingless (Wg) is expressed in the dorsal region of the larval antennal disc (Figure 1C, top left). Because the cells of imaginal discs maintain cell memories through development of the discs (Alexandre et al. 2014), we used the expression domains of these three genes as landmarks to examine the transformation of the antennal disc at eight early pupal stages [from 0 h to 36 h after puparium formation (APF)] (Figure 1C). We found that the primordium of maxillary palp started to dissociate from the antennal disc at about 4 h APF and associated with the developing proboscis prior to 8 h APF (Figure S1A). During this period, the antennal disc also dissociated from the eye disc (Figure S1A). The primordium of arista (arrows in Figure 1C) in the ventral antennal disc began to move anteriorly and dorsally at 8 h APF and eventually reached the lateral region by 12 h APF, corresponding to the hypothesized 90° rotation. This movement occurred along with the ventral extension of midline (*ptc-GAL4*-positive region), followed by the ventral rotation of the Hh-positive posterior compartment. Of note, the 90° rotation of the antennal disc occurred when the head cuticle underwent expansion. By 12 h APF, the time when head eversion occurs, the posterior compartment became ventral, and the dorsal Wg-positive compartment remained in the dorsal region of the developing antenna. The second and third segments of the antennal disc and the base of arista underwent extrusion between 12 h to 16 h APF and was followed by lateral extrusion of the arista primordium at 16 h to 18 h APF. Meanwhile, the Hh-positive compartment in the third segment of developing antenna folded posteriorly along the midline, which resulted in formation of the sac-like structure. By 18 h APF, the sacculus emerged in the posterior region of the antenna. Such tissue transformation resulted in compartmental identities that included an Hh-positive domain in the posterior region, a Ptc-positive midline and a Wg-positive anterior region in the 24 h APF antennae (Figure 1C, 1D).

After associating with the developing proboscis by 8 h APF, the primordium of maxillary palp exhibited a Ptc-positive domain (putatively the midline) in the lateral region, Hh domain in the medial region, and Wg expression in the dorsal region (Figure S1B). The tissue likely underwent folding of the ventral region between 16 h and 18 h APF as the majority of the Hh-positive domain was present in the posterior and Wg-positive regions of the dorsal-anterior developing maxillary palp at 24 h APF (Figure S1B).

Remarkably, the anterior-posterior axis of the antennal disc was initially opposite to that of the body axis at 3rd instar larvae. However, by the end of tissue transformation, the anteroposterior or dorsoventral axes of the antenna were consistent with the global body axes. These observations suggest that the imaginal disc establishes its compartment identities at early stages based on its final orientation in the body plan.

### Morphogens occupy different but partially overlapping domains of antennal discs

In addition to their functions in tissue patterning, morphogens also contribute to neural development (Tiberi et al. 2012). We previously demonstrated that Hh in the peripheral antennal discs/antennae coordinates axon targeting for a subset of ORNs (Chou et al. 2010). Along the same lines, we hypothesized that different morphogens may also occupy different domains of the developing antennal disc/antennae during tissue transformation and later coordinate axon targeting of different subsets of ORNs. In this way, the final coarse compartment map in the periphery after tissue transformation should be coordinated with the axon targeting of ORNs spatially and temporally via combinatorial morphogen codes.

The *Drosophila* genome carries one *hedgehog* (*hh*) gene, seven Transforming growth factor-β family genes [*Activin β* (*Act β*), *dawdle* (*daw*), *decapentaplegic* (*dpp*), *glass bottom boat* (*gbb*), *marverick* (*mav*), *myoglianin* (*myo*), and *screw* (*scw*)], seven Wnt family genes [*wingless* (*wg*), Wnt oncogene analog 2, 4, 5, 6 and 10 (*Wnt 2*, *Wnt 4*, *Wnt 5*, *Wnt 6*, *Wnt 10*), and *wnt inhibitor of Dorsal* (*wntD*)], four Epidermal growth factor receptor agonists [*gurken* (*grk*), *Keren* (*Krn*), *spitz* (*spi*), and *vein* (*vn*)] and three genes encoding Fibroblast growth factor ligands [*branchless* (*bnl*), *pyramus* (*pyr*) and *thisbe* (*ths*)]. We first examined a published single-cell (sc) RNA-seq dataset (McLaughlin et al. 2021) and extracted the expression levels of all annotated morphogens in developing ORNs at when they are targeting to the antennal lobe (24 h APF) and matching with cognate PNs (42 h APF) (Figure 2). The results suggested that morphogens were likely differentially expressed in different populations of developing ORNs. Since the scRNA seq data did not cover all types of ORNs not neighboring supporting cells, we next examined the expression patterns of these ligands in the antennal discs of late 3rd instar larvae and in 24 h APF antennae (Figure 3). Limited by the availability of specific antibodies, we examined some of these ligands through GFP protein trap or gene reports, the latter reflects the gene expressions instead of protein. We found that ligands of each family are expressed in different domains of the antennal disc and 24 h APF antennae. Interestingly, some other ligands, such as Vn, were not expressed in the antennal disc but were later expressed in the developing antennae at 24 h APF. Collectively, these results suggested that morphogens of Hh, TGFβ, Wnt, EGF and FGF families are expressed in the antennal disc and developing antennae; however, the expression of particular ligands changes at different developmental stages.

**Figure 2.**
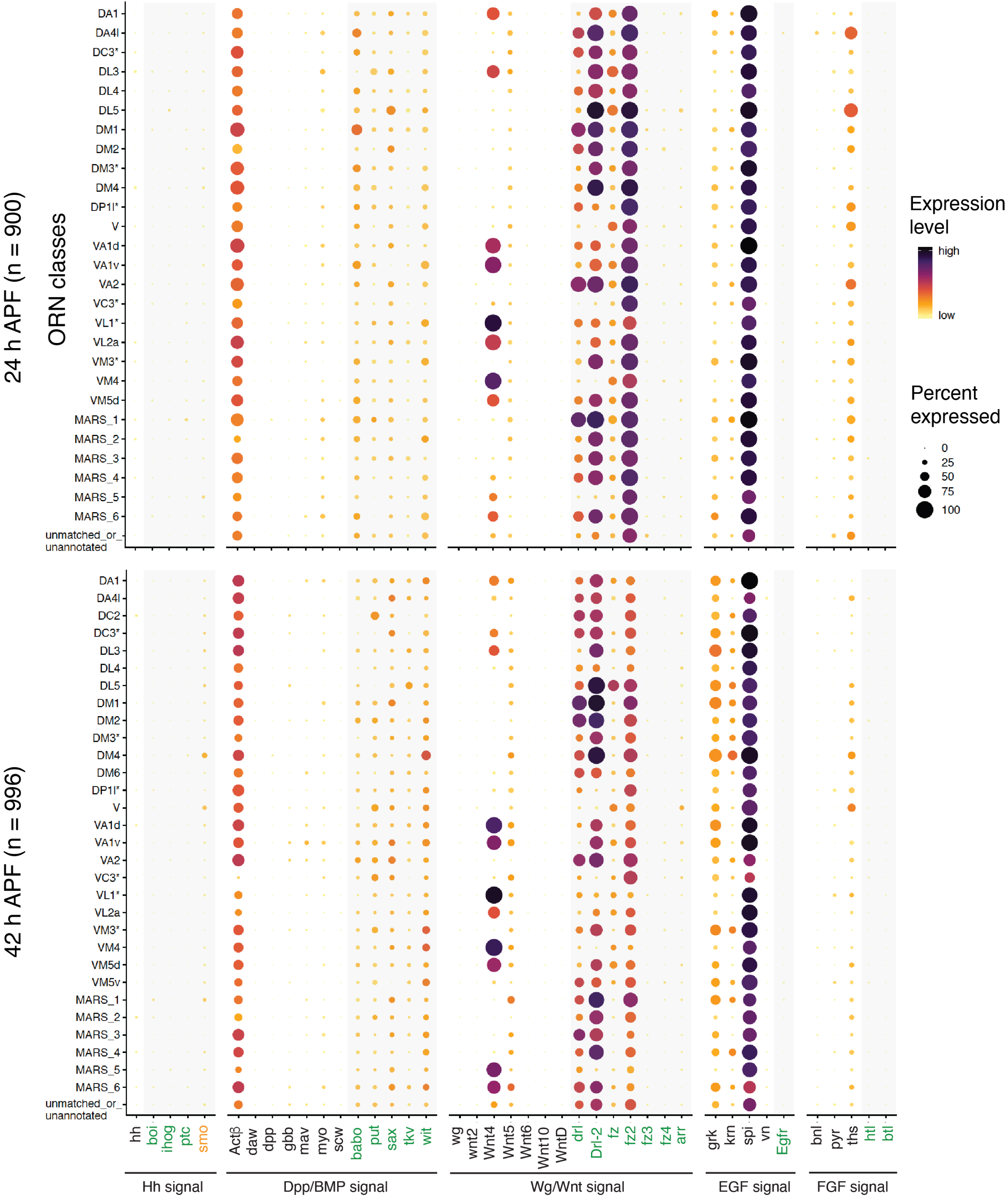
Single-cell and single-nuclei RNA-seq analyses of morphogen and morphogen receptor expression levels in ORNs. Dot plots summarizing the expression of ligands and receptors of Hh, TGFβ, Wg/Wnt, EGF and FGF signaling pathways in different ORN types at 24 h APF, 42 h APF and adult stage.

**Figure 3.**
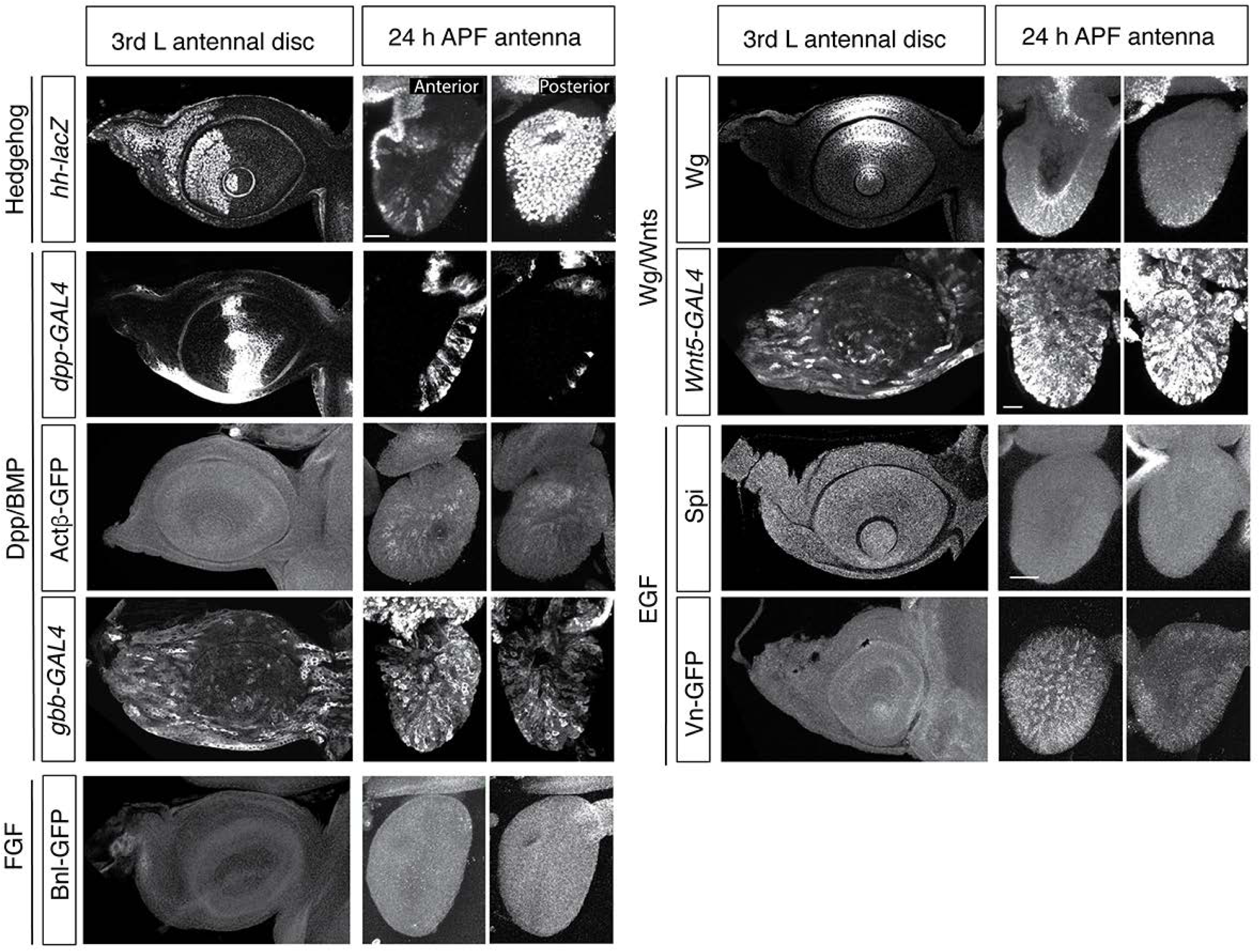
Differential expression patterns of morphogen ligands in antennal discs and 24 h APF antennae. Representative confocal images of late 3rd instar larval antennal discs and 24 h APF antennae (with partial projections of anterior sections and posterior sections). The expression patterns of morphogens were examined either by expression of reporters (*hh-lacZ*, *dpp-GAL4*, *Gbb-GAL4*, and *Wnt5-GAL4*) or antibody staining (Actβ-GFP, Bnl-GFP, Wg, Spi, and Vn-GFP).

### Morphogens likely coordinate the spatial information of ORNs during tissue transformation

Neurogenesis and specification of ORNs occur from 0 h to about 20 h APF (Endo et al. 2007; Brochtrup and Hummel 2011; Chai et al. 2019; Endo et al. 2011). To gain a better understanding of the timing for ORN soma emergence and their axon innervation, we used pan-ORN driver *peb-GAL4* to visualize ORNs from late 3^rd^ instar larva to mid-pupal stages (Figure 4). A subset of ORNs began to extend their axons at 15 h APF, and all ORN axons fasciculated into two axon bundles by 18 h APF (Figure 4). The axons of ORNs had arrived in the developing antennal lobe at 18 h APF and began to innervate the AL at around 24 h APF (Jefferis et al. 2004; Jhaveri et al. 2000) (Figure 4). Cell bodies of ORNs are housed in three different types of sensilla (basiconic, trichoid and coeloconic), which are distributed among different domains of the antenna (Shanbhag et al. 1999). The glomerular targeting by specific ORN classes in each distinct sensillar type was restricted to a roughly unique region of the antennal lobe, demonstrating coordination of the peripheral sensilla map and central ORN targeting/glomerular map (Couto et al. 2005). Such coordination remains to some extent when comparing the soma distributions of different types of ORNs in the antenna and their glomerular targets in the antennal lobe (Chai et al. 2019). It was also previously shown that positional patterns and cell motility may contribute to the dispersed patterns of distinct types of ORN somas in the antenna (Song et al. 2012). The possible “driving force” of such cell motility remains unclear.

**Figure 4.**
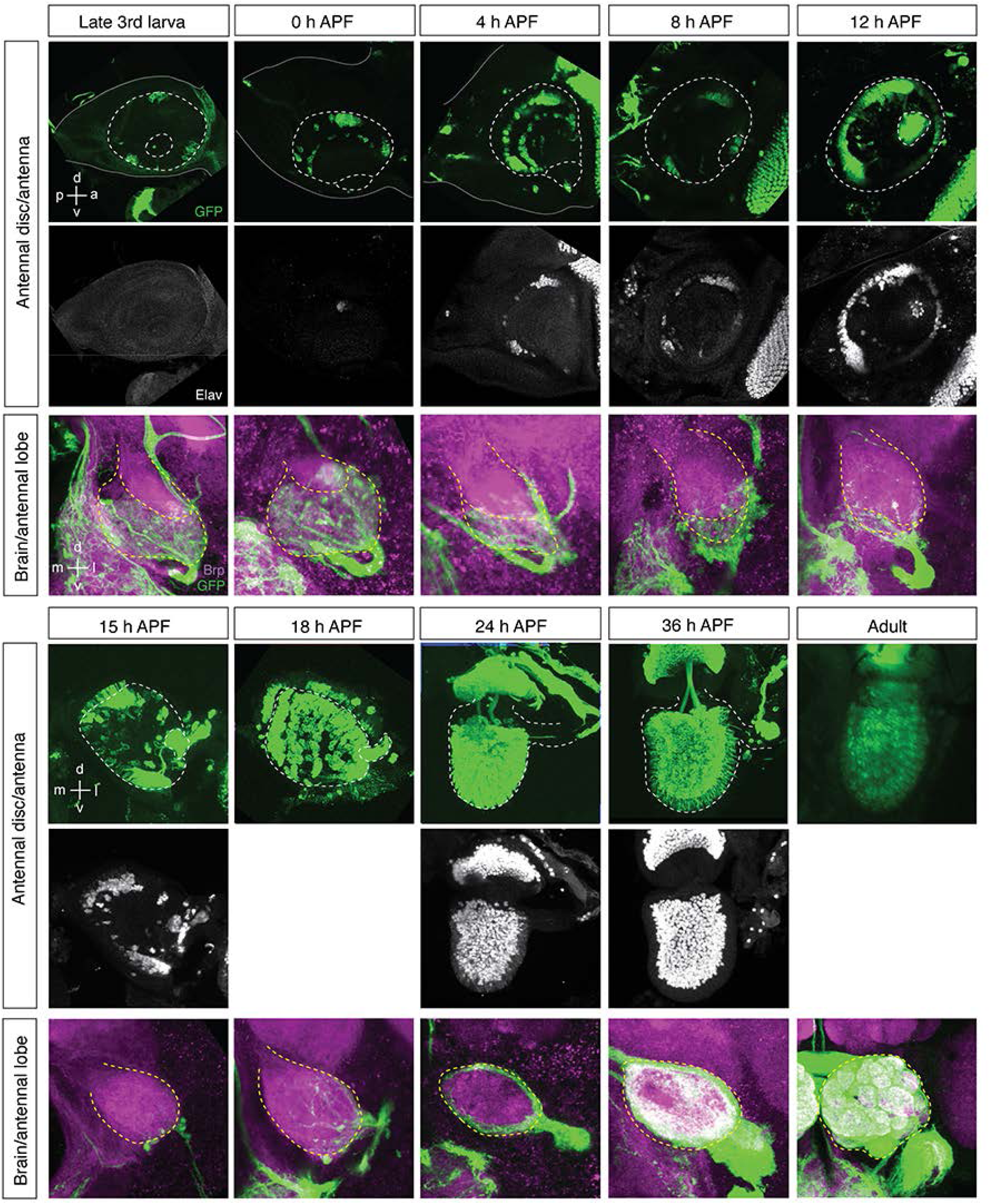
Axon targeting of all ORNs during development. Late 3rd instar larval (3L) antennal discs and pupal antennae were stained for *peb-Gal4*-driven mCD8GFP (green) and pan-neuronal marker Elav (magenta). Segment boundaries were determined by DAPI staining. Yellow solid lines, white solid lines and white dashed lines mark the contours of antennal discs, 2nd segments and 3rd segments, respectively. Arrowheads point to aristae. *peb-Gal4* is not expressed in the 3rd segment of larval antennal discs. At 0-4 h APF, *peb-Gal4* is expressed in a few cells in the 3rd segment. At 8-12 h APF, *peb-Gal4* expression is somewhat reduced. At 18 h APF and later, *peb-Gal4* is strongly expressed in all antennal cells. All images were acquired through confocal microscopy, except the one showing adult antennae, which was acquired from unfixed samples through fluorescence microscopy. Scale bars, 50 μm.

We noted that the events of ORN neurogenesis, specification and initiation of axon targeting all occur during the period of dramatic tissue transformation from the antennal disc to the antenna. This timing means that the cellular events of ORNs are somehow temporally coordinated with the tissue transformation of antennal disc to antenna. Therefore, we next asked how such temporal coordination may be achieved and whether it would partially contribute to the spatial coordination between ORN soma positions in the antenna and axonal targeting in the antennal lobe. Two lines of evidence suggest that morphogens are likely candidates to coordinate ORN cellular events and tissue transformation. First, our tissue transformation results showed that cells expressing the same morphogen moved together during antennal disc transformation from 0 h to 24 h APF (Figure 1C). Second, Hh signaling has been shown to coordinate the peripheral map and the central axonal targeting map (Chou et al. 2010). If morphogens are indeed mediators of this coordination, we would expect that the corresponding receptors of these morphogens are also expressed in different subsets of ORNs at 24 h APF during axon targeting and even later when cognate ORNs and projection neurons (PNs) undergo matching. We therefore reanalyzed single-cell (sc) RNA-seq and single-nucleus (sn) RNA-seq data and found different expression levels of Hh receptor, Type I and Type II TGFβ receptors, and Wnt receptors in different types of ORNs at 24 h and 42 h APF (Figure 2). Interestingly, although the corresponding ligand expression levels were high (*spi*, *grk*, and *ths*), the expression levels of some receptors (*EGFR*, *htl* and *btl*) in ORNs were low. These results imply that morphogens likely coordinate the spatial distributions of ORN somas during tissue transformation. To further verify this idea, we next investigated the possible involvement of morphogens in ORN axon targeting.

### PNs may provide morphogens for ORNs

Since we considered morphogen signals to be likely candidates for temporal coordination of tissue transformation and cellular events of ORNs, including axon targeting, we expected that we would be able to identify the sources of morphogens in developing antennal lobes and observe differential mistargeting phenotypes of distinct subsets of ORNs in morphogen receptor mutants. Indeed, we observed the expressions of some morphogens around 24 h APF antennal lobes (Figure 5A).

**Figure 5.**
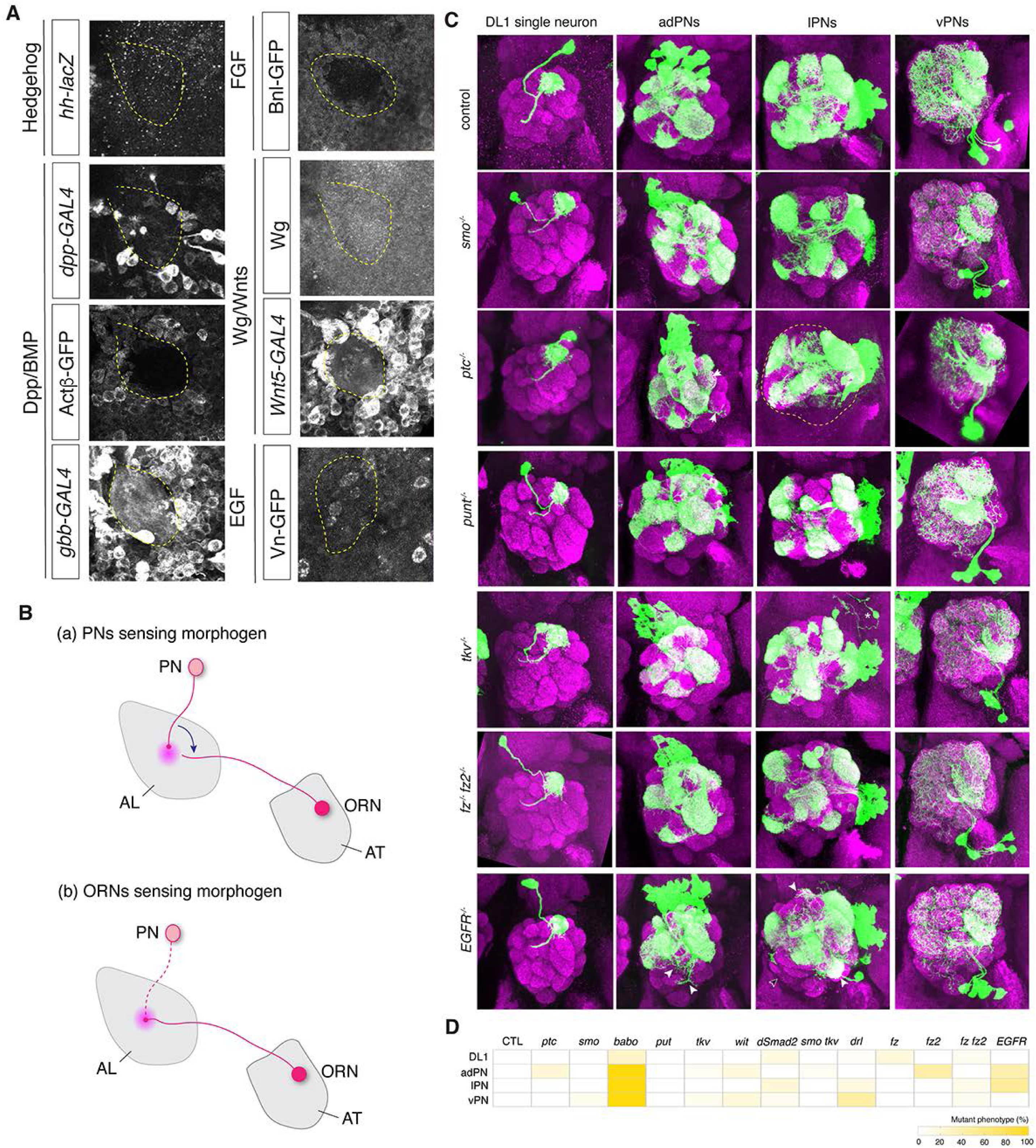
Projection neurons express morphogens but do not require morphogens for dendrite development. (A) Representative confocal images of 24 h APF antennal lobe. The expression patterns of morphogens were examined either by expression of reporters (*hh-lacZ*, *dpp-GAL4*, *Gbb-GAL4*, and *Wnt5-GAL4*) or antibody staining (Actβ-GFP, Bnl-GFP, Wg, and Vn-GFP). (B) Schematic diagram of two possible mechanisms for morphogens signals (light pink) in the antennal lobe (AL) in ORN axon targeting. Morphogen receptors (deep pink) are expressed in different subsets of PNs and ORNs. AT, antennae. See text for the details. (C) Representative confocal images of DL1 PN single cell clones, adPNs neuroblast clones, lPN neuroblast clones and vPN neuroblast clones carrying different morphogen receptor mutations. White arrowheads indicate abnormal dendrite extensions. (D) Heatmap showing the degrees of axonal mistargeting phenotypes.

As soon as flies enter the pupal stage, their PNs undergo pruning and dendrite re-extension at around 6 h APF and form a coarse glomerular map at around 24 h APF (Jefferis et al. 2004). The axons of ORNs in the antennae (AT ORNs) first arrive at the AL at 18 h APF and innervate the developing AL where they find cognate PNs later (Komiyama et al. 2007; Zhu et al. 2006; Hong et al. 2009; Lattemann et al. 2007). The innervation of AT ORNs in the antennal lobe constrains the target choice of late-arriving axons from ORNs in the maxillary palps (MP ORNs) (Sweeney et al. 2007). Although we cannot exclude that other neurons around the developing antennal lobe, such as local interneurons (Liou et al. 2018) and multi-glomerular projection neurons (Tanaka et al. 2012; Lin et al. 2012), may provide ligands for ORN axon targeting, PNs are the most likely candidates to provide morphogens to the ORNs expressing different morphogen receptors. To test this idea, we reanalyzed published scRNA-seq data (Xie et al. 2021), systematically exploring the expressions of all morphogens and receptors (Figure S4). Indeed, PNs differently expressed Hh, Actβ, Mav, Gbb, Wnt5, Spi, Grk, Pyr and Ths at 24 h and 42 h APF. However, we noticed that some PNs also expressed Hh signal effector Smo, all five TGFβ receptors, and four Wnt receptors (Drl, Drl2, Fz and Fz2) (Figure S4). These results raised two possible mechanisms for morphogen-guided ORN axon targeting. First, developing PNs may receive different morphogen signals to build their targeting map in the antennal lobe, which in turn guides ORN axons to the cognate glomeruli (Figure 5B, top). Second, ORNs axons sense morphogens for their targeting (Figure 5B, bottom). To assess the first possible mechanism, we asked whether the morphogen receptor mutations would lead to PN dendrite defects (Figure 5C, D, Figure S3, Table 1). Except for a previously reported pruning defect in *bobo* mutants (Marin et al. 2005), PNs did not appear to typically rely on morphogen signals for their dendrite development. Thus, PNs may be at least one of the cellular sources of morphogen ligands for ORN axon targeting and the expressed morphogen signals are not utilized for PN dendrite development.

**Table 1.**
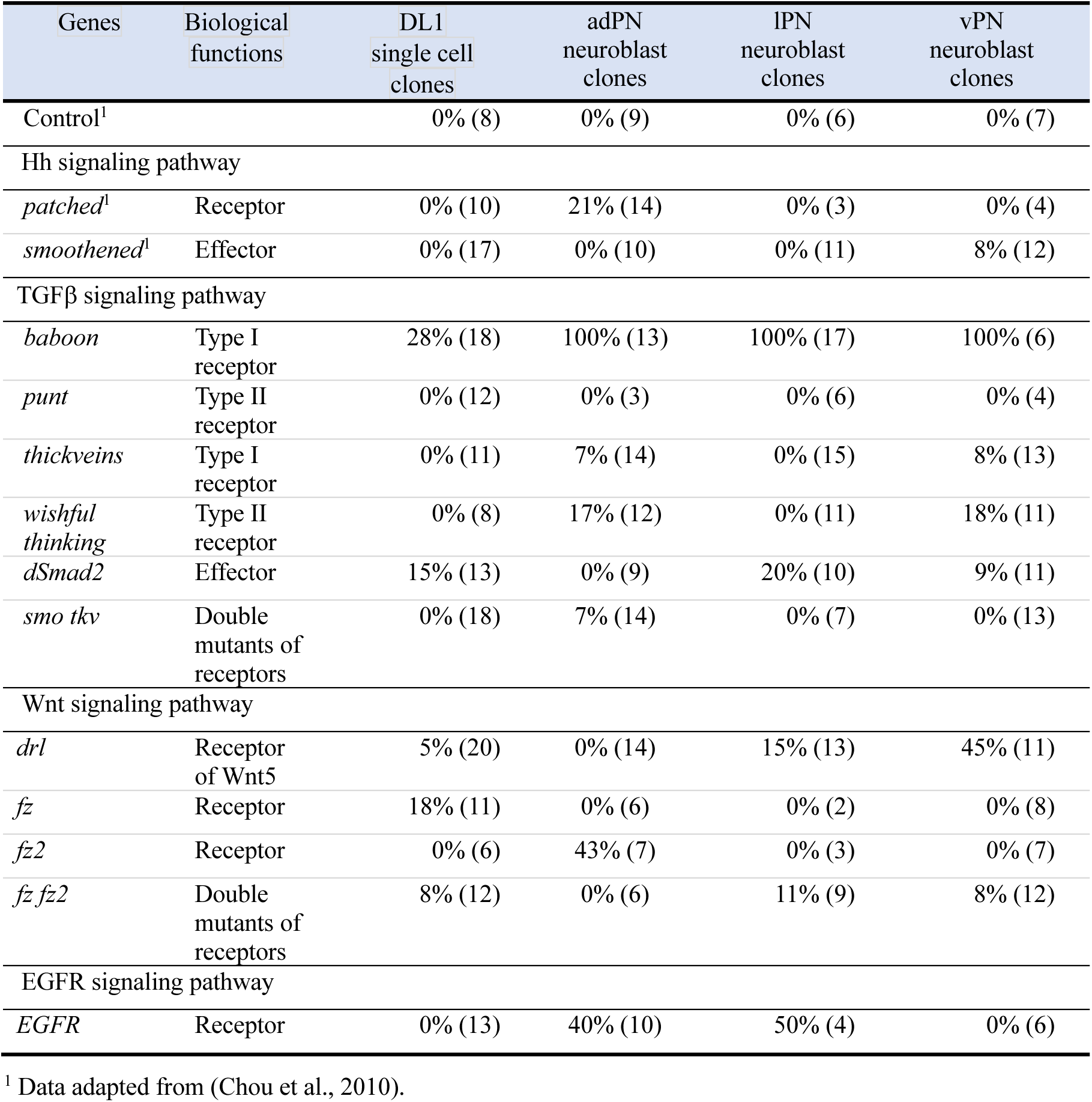
Projection neuron receptor and effector mutant phenotypes.

### Differential requirement of morphogen signals for ORN axon targeting

To test the second possible mechanism that morphogen signals are required for ORN axon targeting (Figure 5B, bottom), we would expect that ORNs carrying receptor mutants should show mistargeting phenotypes. In line with this idea, *smo* mutant clones of distinct ORN types demonstrated differential requirements for Hh signaling for axon targeting (Chou et al. 2010). Similarly, the Wnt 5 receptors, Drl and Drl 2, have been shown to exert opposite effects on ORN axon targeting (Sakurai et al. 2009). FGF signals from LNs to FGF receptors on wrapping glia are also known to be required for PN and ORN matching (Wu et al. 2017). In addition, both FGF receptors, *htl* and *btl*, were absent or barely expressed in analyzed ORNs at 24 h and 42 h APF (Figure 2). Therefore, we focused on evaluating the requirements for TGFβ receptor Tkv, Wg/Wnt receptors Fz and Fz2, and EGFR in ORN axon targeting.

Morphogens at early stages are required for the growth of imaginal discs (Lebreton et al. 2008; Alexandre et al. 2014). To avoid any possible mutant clones in the brain and to prevent secondary effects due to defects in early antennal disc growth, we used ey-FLP to introduce mutant clones of morphogen receptors in antennae and maxillary palps. We then examined axonal targeting of mutant ORNs, excluding a small subset of flies that carried obvious antenna and/or maxillary palp malformations from the analysis (Figure 6, Figure S4, Table 2).

**Table 2.** ey-FLP-mediated ORN receptor mutant phenotypes.

| ORN class | Glomerular target(s)<sup>1</sup> | AT/MP<sup>2</sup> | WT (40A)<sup>3</sup> | *tkv*<sup>-/-</sup> | *EGFR*<sup>-/-</sup> | *fz*<sup>-/-</sup> *fz2*<sup>-/-</sup> | *smo*<sup>-/-3</sup> | N<sup>4</sup><br>(on/off) |
| --- | --- | --- | --- | --- | --- | --- | --- | --- |
| Or9a | VM3 | AT | 15% (26) | n.d.<sup>5</sup> | n.d. | n.d. | 26% (19) | n.t.<sup>4</sup> |
| Or10a | DL1 | AT | 8% (24) | 0% (10) | 77% (13) | n.d. | 8% (26) | On |
| Or13a | DC2 | AT | 4% (28) | n.d. | n.d. | n.d. | 28% (25) | n.t. |
| Or22a | DM2 | AT | 0% (20) | 0% (22) | 0% (18) | 5% (22) | 4% (24) | On |
| Or23a | DA3 (DC3) | AT | 5% (21) | 54% (13) | 0% (16) | n.i (19) | 0% (28) | On |
| Or42a | VM7 (V, VL2p) | MP | 4% (24) | 40% (15) | 32% (22) | 22% (9) | 7% (28) | Off |
| Or43a | DA4l | AT | 9% (22) | 79% (19) | 8% (13) | 0% (18) | 72% (25) | On |
| Or43b | VM2 (DM5) | AT | 0% (9) | n.d. | n.d. | n.d. | 13% (15) | Off |
| Or46a | VA7l | MP | 10% (20) | n.i. (28) | 0% (23) | n.d. | 25% (44)<sup>6</sup> | Off |
| Or47a | DM3 | AT | 5% (21) | 35% (23) | 0% (20) | 70% (27) | 3% (29) | On |
| Or47b | VA1v | AT | 0% (21) | 0% (18) | 5% (20) | 75% (20) | 0% (20) | Off |
| Or49a | DL4<sup>7</sup> | AT | 0% (22) | n.d. | n.d. | 0% (2)<sup>7</sup> | 0% (21) | On |
| Or49b | VA5 | AT | 8% (13) | n.d. | n.d. | n.d. | 19% (36) | n.t. |
| Or56a | DA2 | AT | 10% (40) | n.d. | n.d. | n.d. | 35% (40) | Off |
| Or59c | 1 (VM7) | MP | 13% (16) | 4% (28) | 0% (10) | 100% (16) | 55% (22) | Off |
| Or67a | DM6<sup>7</sup> | AT | 0% (22) | n.d. | n.d. | 0% (2)<sup>7</sup> | 43% (21) | Off |
| Or67b | VA3 | AT | 14% (21) | 5% (21) | 0% (21) | 33% (9) | 5% (22) | Off |
| Or71a | VC2 | MP | 6% (17) | 0% (5) | 20% (10) | 40% (15) | 0% (26)<sup>6</sup> | On |
| Or83c | DC3 (VA6) | AT | 5% (21) | n.i. (23) | 0% (8) | 13% (8) | 38% (21) | Off |
| Or85e | VC1 | MP | 0% (20) | 0% (21) | 38% (21) | 60% (20) | 50% (20) | On |
| Or88a | VA1d | AT | 0% (20) | 35% (23) | 0% (19) | n.d. | 0% (20) | n.t. |
| Or92a | VA2 | AT | 0% (20) | 65% (20) | 0% (20) | 54% (13) | 5% (20) | Off |
| Gr21a | V | AT | 9% (22) | 42% (26) | n.d. | 24% (17) | 0% (25) | On |

<sup>1</sup>Glomeruli targeted by ectopically labeled ORNs are indicated in the parentheses and were not included in phenotype quantification.
<sup>2</sup> AT, ORN soma in antennae; MP, ORN soma in maxillary palps.
<sup>3</sup> Data adapted from (Chou et al., 2010), except for the results of Or49a and Or67a.
<sup>4</sup> Data adapted from (Endo et al., 2007).
<sup>5</sup> n.d., not determined; n.i., none of examined brains were innervated by GFP-positive axons; n.t., not determined due to ambiguous results.
<sup>6</sup> Data are from hs-FLP-mediated mutant clones.
<sup>7</sup> ORNs were labeled by AM29-GAL4. Among 11 analyzed brains, only two brains had GAL4 expression.

By systematically examining the axonal targeting phenotypes of 15 ORN classes carrying *tkv*, *fz fz2* or *EGFR* mutations, we identified differential requirements of morphogen signals to distinct classes of ORNs. For instance, *tkv* mutant Or43a ORNs mistargeted to other glomeruli while *EGFR* or *fz fz2* mutant Or43a ORNs did not show abnormal targeting (Figure 6A, top row). Or47b ORNs exhibited mistargeting phenotypes in all three mutant backgrounds (Figure 6A, second row). In contrast, Or92a ORN axons showed different mistargeting phenotypes when they were *tkv* or *EGFR* mutants but no phenotype with *fz fz2* mutation (Figure 6A bottom row). ORNs from maxillary palps also showed differential requirements for these three morphogen signals (Figure 6A, third row, Figure S4). These results demonstrated that ORNs employ combinatorial morphogen codes for their axonal targeting (Figure 6B).

## Discussion

### Temporal coordination of tissue transformation and sensory neuron axonal targeting

During pupal development, the two-dimensional antennal disc undergoes a series of tissue transformations, including dissociation from the eye disc and maxillary palp primordium, transformation of two-dimensional antennal disc to three-dimensional antennae, extrusion of arista, formation of sacculus, and migration and integration of maxillary palp primordium to the labial disc (Figure 7, top). These events are tightly coordinated with neurogenesis of ORNs in the antennal disc, the acquisition of sensory neural fate and sensillum fate, and the axonal targeting of sensory neurons from the antenna to the central brain (Figure 7. Middle). Such coordination is likely mediated by dual tissue patterning and axon guidance functions of morphogens, which act together as combinatorial codes to guide the development of antennae globally and individual sensory neurons locally.

**Figure 6.**
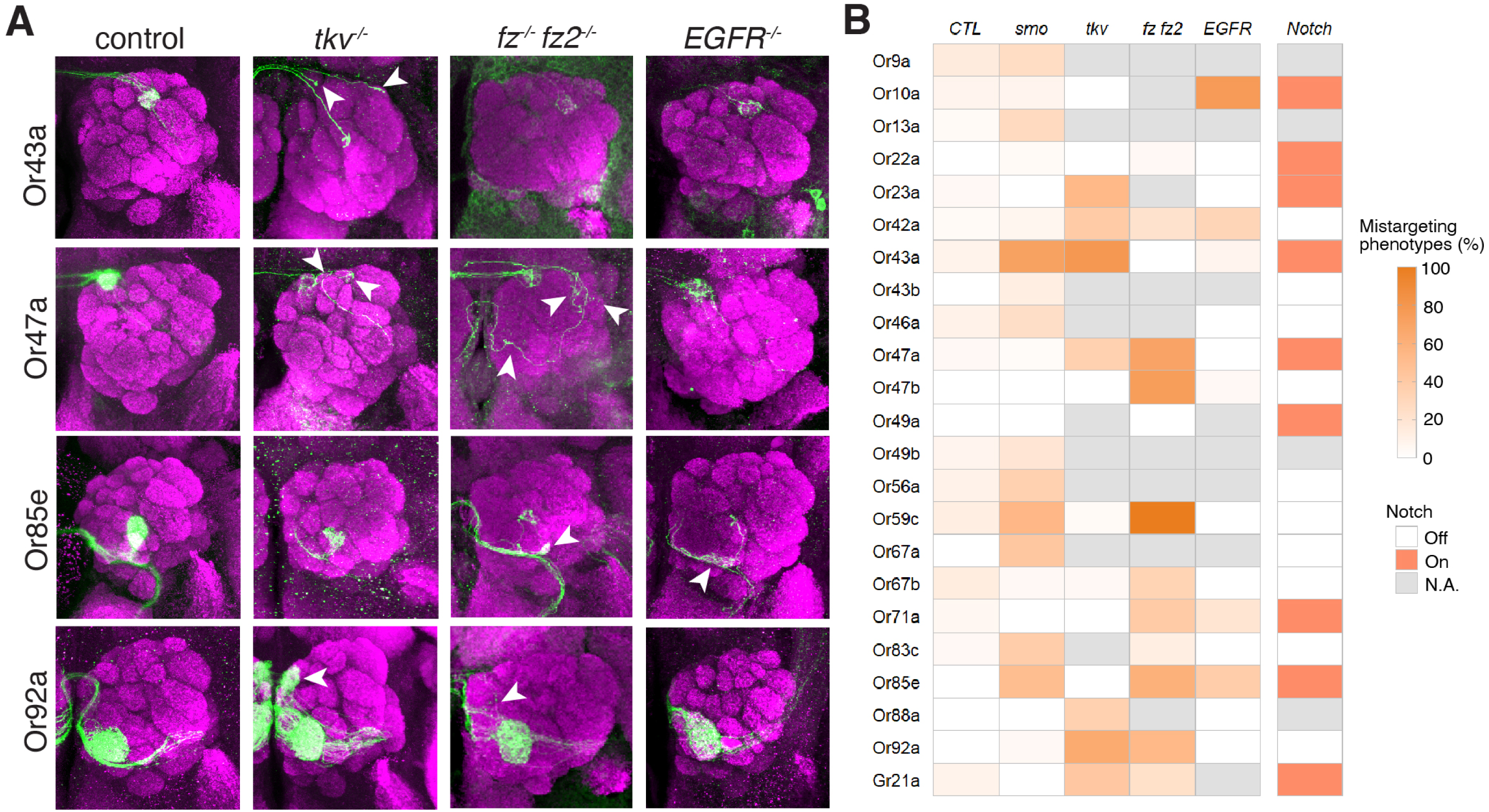
Combinatorial codes of morphogens regulating ORN axon targeting. (A) Representative confocal images showing axonal targeting of different ORN classes in the antennal lobes. White arrowheads indicate mistargeted axons. (B) Heatmap showing the degrees of axonal mistargeting phenotypes.

**Figure 7.**
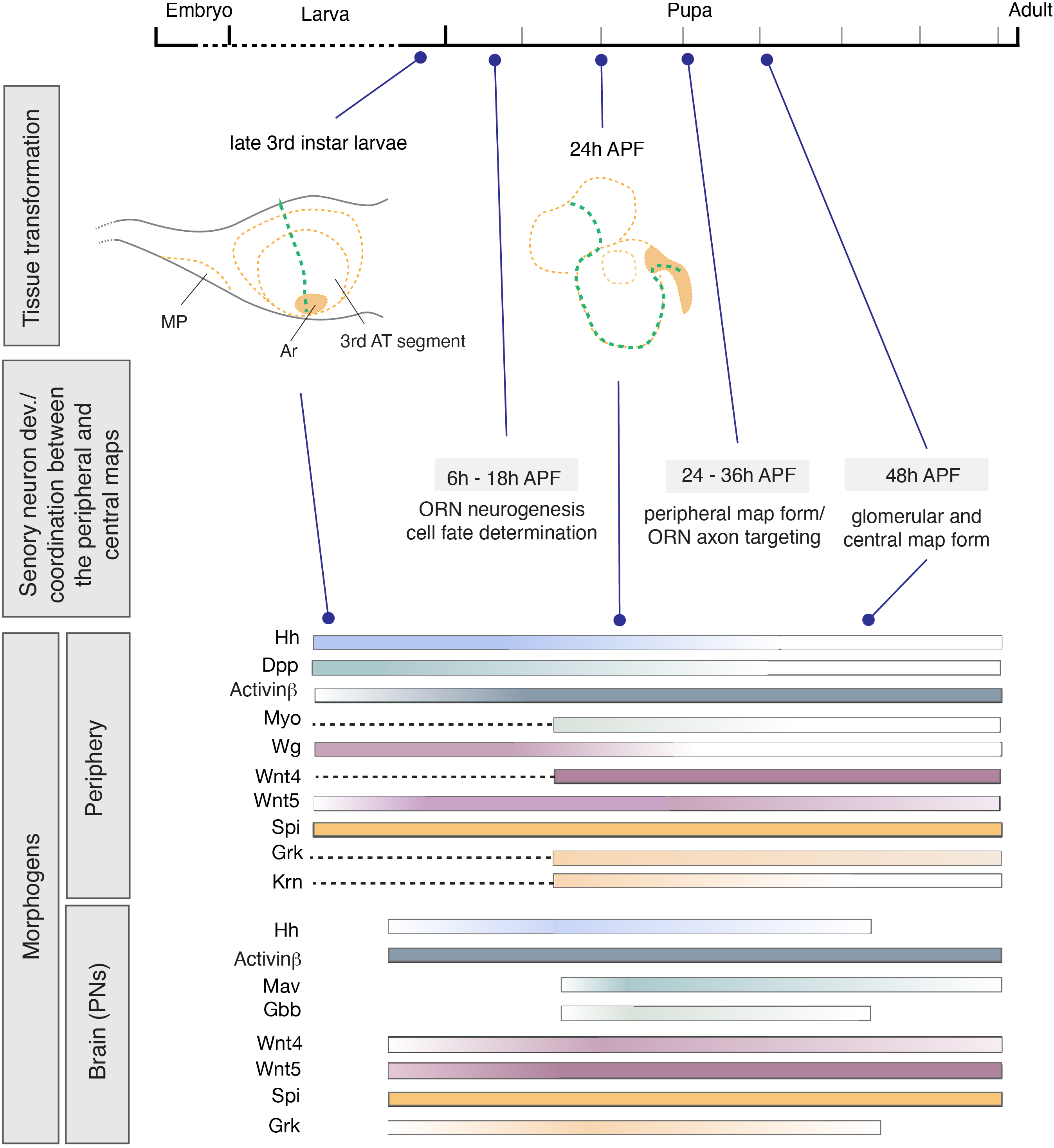
Temporal coordination of tissue transformation and ORN axon targeting. (Top) The morphological changes of antennal disc to antennae from late 3^rd^ instar larval stage to 24 h APF pupal stage. See text for details. (Middle) Corresponding timings of ORN cell fate determination, ORN axon targeting and final coordination between the peripheral maps (sensilla) and the central map (ORN glomerular targets). (Bottom) Temporal switch of morphogens in the periphery (antennal discs/antennae) and in the brain.

Interestingly, the patterning of larval antennal discs occurs such that the disc aligns its tissue axes with the final adult body axes of adult and not the global body axes of larva. Another interesting question is related to the identity of the driving force for tissue dissociation. This driving force may involve a local mechanical force within the antennal disc or “dragging” by the head cuticle as it expands. It also remains a mystery how the primordium of maxillary palp finds its way to meet and integrate with the labial disc. Future work should focus on understanding the mechanisms underlying these events.

### Two strategies of coordination by morphogens

Although the availability of specific antibodies or gene reports of all ligands limits the observation of morphogens profiled through scRNA-seq analyses of developing ORNs, both sets of analyses demonstrated differential expressions of ligands within the same families and across different developmental stages (Figure 2, Figure 3). Based on these observations and receptor mutans (Figure 6, Figure S4), we found two molecularly distinct strategies of morphogens are likely employed to coordinate the tissue transformation and ORN axon targeting: (1) a two-step strategy and (2) switching ligands. An example of the two-step strategy is Hh (Chou et al. 2010), the only ligand of the Hh signaling pathway in *Drosophila*. At the patterning stage, Hh from the posterior compartment establishes the anterior-posterior boundary through binding to Patched in the midline cells (Figure 7, bottom). This action in turn causes cells, including future ORNs, in the antennal disc to be either Ptc-negative cells or Ptc-positive cells. The Ptc-positive ORNs no longer respond to brain-derived Hh during their axon targeting (Figure 7, bottom).

In contrast to the Hh signal, TGFβ, Wnt and EGF signals exhibit ligand switch strategies to coordinate antennal disc patterning and ORN axon targeting. For instance, the TGFβ ligand Dpp is expressed in the antennal disc and functions to pattern the ventromedial region of the antennal disc, while another TGFβ ligand, Actβ, is increased at the early pupal stage both in the peripheral antennal disc and in the brain (Figure 7, bottom). In addition to Actβ, Mav and Gbb are expressed in the central brain during ORN axon targeting stage. Nevertheless, how such a switch is controlled remains unclear.

### Coordination between the peripheral map and the central map

A coarse correlation between the sensory neuron distribution in mouse olfactory epithelium and axon targeting to glomeruli in olfactory bulbs has been reported (Miyamichi et al. 2005; Zhu et al. 2022). This phenomenon also seems to be evolutionary conserved in *Drosophila*. At the sensillar level, the distribution of sensory somas in the antennae and maxillary palps (the peripheral map) exhibits a coarse correlation with the targeted glomeruli in the antennal lobe (the central map) (Couto et al. 2005; Chou et al. 2010). However, this correlation of the peripheral map and central map in *Drosophila* becomes weaker (but remains significant) at the sensory neuron level in the antennae (Chai et al. 2019). Local “dispersion” of sensilla during antennal disc transformation was observed, accompanied by the switching-on of a posterior marker Engrailed in a few anterior cells near by the antero-posterior boundary (Song et al. 2012). On top of this observation, the Patched-positive cells originating in the anterior-posterior boundary become intercalated with Patched-negative cells in the anterior-posterior boundary at 24 h APF antennae (Figure 1C) (Song et al. 2012; Chou et al. 2010). The dramatic tissue transformation described herein may be one of the driving forces leading to such sensillar dispersion.

### Restricted requirement of morphogen signals for sensory neuron development

We found that combinatorial codes are used by different ORN classes for axon guidance. Although we cannot rule out that other neurons or glia around the antennal lobe provide morphogens to guide incoming ORN axons, scRNAseq analyses of PNs demonstrated that different PNs express distinct sets of morphogens. For instance, at 24 h APF, DA4l PNs express Actβ, Mav, Wnt5, Grk, Spi and Vn; meanwhile, VA1v PNs express Actβ, Wnt5, Grk and Spi. On the other hand, Actβ, Wnt5, Grk and Spi are expressed in all examined PNs, but the expression levels vary across different types of PNs (Figure S2).

At 24 h APF, DA4l ORNs express tkv (weakly), Drl, Drl2 and Fz2, but did not express EGFR (Figure 2). MARCM analyses of OR43a ORNs (to DA4l) further revealed mistargeting phenotypes with tkv, fz fz2 mutations but not EGFR mutation (Figure 6B). Or92a ORNs (to VA2) are Tkv-positive, Drl/Drl2-positive, Fz/Fz2-positive but EGFR-negative, while *fz fz2* mutant ORNs normally target the DA2 glomerulus (Figure 6A). This finding is not surprising because molecules that promote cognate PNs dendrite and ORN axon matching at approximately 30 h to 42 h APF may partially compensate for the phenotypes (Zhu and Luo 2004; Zhu et al. 2006; Hong et al. 2009; Sweeney et al. 2007; Ward et al. 2015; Hong et al. 2012; Xu et al. 2024). Remarkably, PNs with examined morphogen receptor mutants did not show obvious dendrite defects. Therefore, we conclude that the morphogen codes are mainly restricted to peripheral sensory neurons and are not employed by second order PNs in the brain.

## Author Contributions

Y.H.C. designed and supervised the project. All experiments were performed and analyzed by S.C.J.C, Y.H.C., and I.L, except the scRNAseq and snRNAseq analyses, which were performed by

T.M.L and C.T.L. Y.H.C., S.C.J.C, and T.M.L wrote manuscript.

## Data and Code Availability

The original, unprocessed confocal images and scripts used in this study will be made available upon reasonable request.

## Acknowledgements

Y.H.C thanks Dr. Liqun Lou for facilitating the project; some preliminary results were initiated in his lab by Y.H.C. We thank Drs. Tetsuya Tabata, Makoto Sato, Henry Y. Sun, Chihiro Hama for sharing flies. Except of requested reagents, stocks and antibodies obtained from the Bloomington Drosophila Stock Center (NIH P40OD018537), KYOTO Drosophila Stock Center in Kyoto Institute of Technology, Japan), and Developmental Studies Hybridoma Bank were used in this study. We are also thankful for help from Taiwan flycore and DNA Sequencing Core Facility. We thank Dr. Marcus Calkins for English editing. This work was supported by Academia Sinica Grand Challenge Program (AS-GC-112-L01), and National Science and Technology Council (NSTC 112-2311-B-001-036-MY3) to Y.-H.C.

---

**Figure S1. The development of maxillary palps during pupal stages.** (A) Confocal image of 8 h APF head cuticle whole mount showing the majority of developing imaginal discs in their relative positions and orientations. R, right; L, left. (B) Confocal images of developing maxillary palps at different pupal stages. Compartments of the maxillary palps are labeled by *hh-lacZ* (red), *ptc-GAL4* driven nuclei-GFP (green), and Wg (blue). Scale bars, 100 μm.

**Figure S2. Single-cell RNA-seq analyses of morphogen and morphogen receptor expression levels in PNs.** Dot plots summarizing the expression of ligands and receptors in the Hh, TGFβ, Wg/Wnt, EGF and FGF signaling pathways in PN types at 0 h APF, 24 h APF, 48 h APF and adult stage.

**Figure S3. Dendrite development of PNs carrying morphogen receptor and effector mutations** Representative confocal images of DL1 PN single cell clones, adPN neuroblast clones, lPN neuroblast clones and vPN neuroblast clones that carried different morphogen receptor mutations. White arrowheads indicate abnormal dendrite extensions.

**Figure S4. Axonal targeting of ORNs carrying morphogen receptor and effector mutations.** Representative confocal images showing axonal targeting of different ORN classes in the antennal lobes. White arrowheads indicate mistargeted axons.

