## Supplementary for "Temporal coordination of tissue transformation, olfactory sensory neural development and central axon projections through morphogens"

### Supplementary Figures

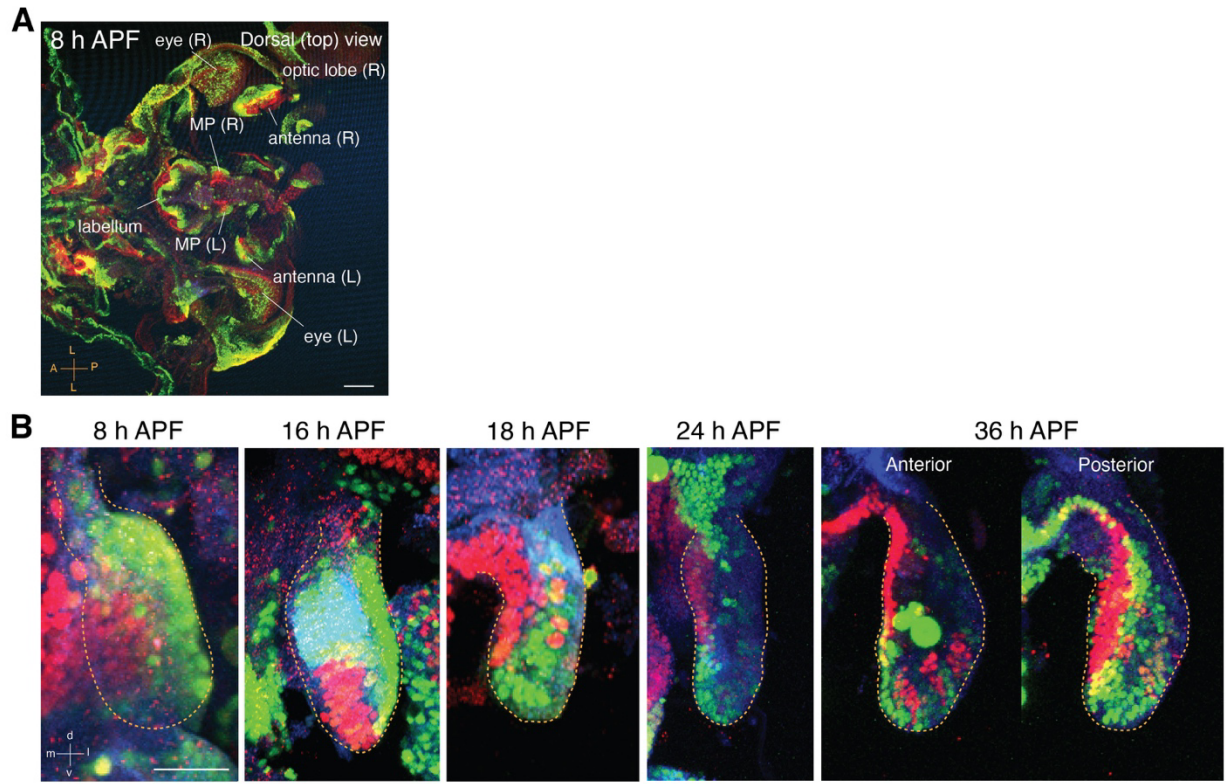

**Figure S1. The development of maxillary palps during pupal stages.**

(A) Confocal image of 8 h APF head cuticle whole mount showing the majority of developing imaginal discs in their relative positions and orientations. R, right; L, left.

(B) Confocal images of developing maxillary palps at different pupal stages. Compartments of the maxillary palps are labeled by *hh-lacZ* (red), *ptc-GAL4* driven nuclei-GFP (green), and Wg (blue). Scale bars, 100 μm.

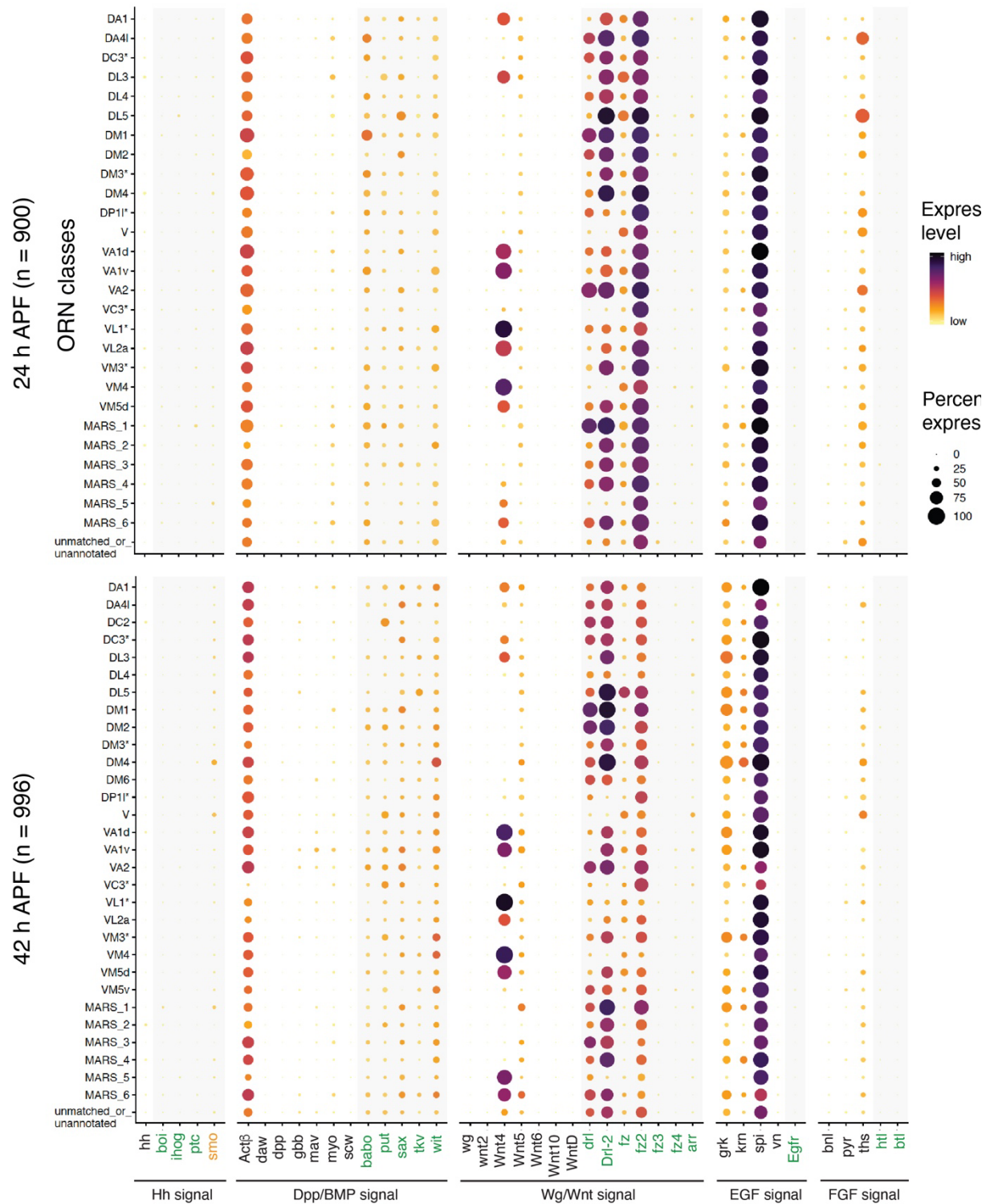

**Figure S2. Single-cell and single-nuclei RNA-seq analyses of morphogen and morphogen receptor expression levels in ORNs.**

Dot plots summarizing the expression of ligands and receptors of Hh, TGFb, Wg/Wnt, EGF and FGF signaling pathways in different ORN types at 24 h APF, 42 h APF and adult stage.

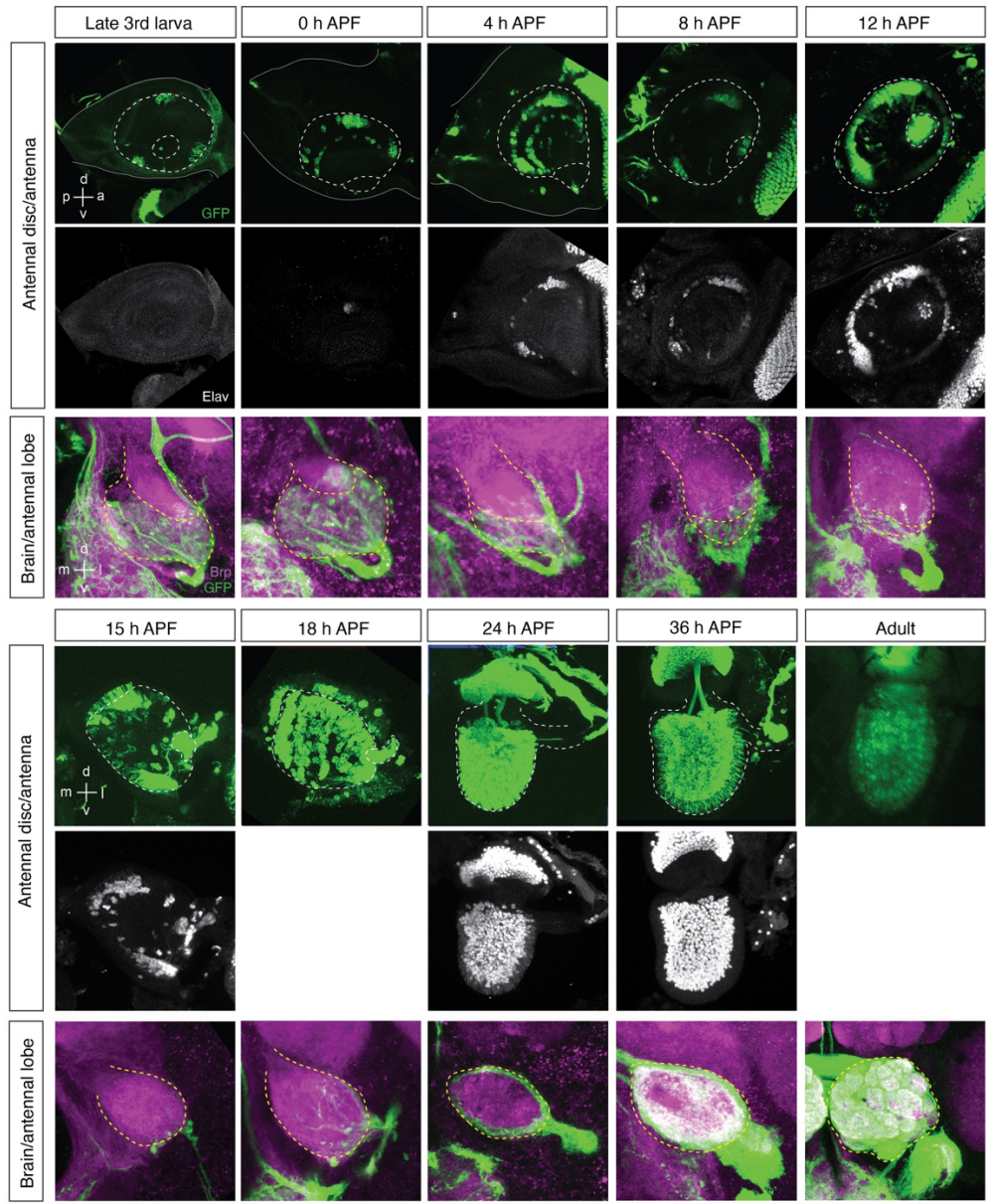

**Figure S3. Axon targeting of all ORNs during development.**

Late 3rd instar larval (3L) antennal discs and pupal antennae were stained for *peb-Gal4*-driven mCD8GFP (green) and pan-neuronal marker Elav (magenta). Segment boundaries were determined by DAPI staining. Yellow solid lines, white solid lines and white dashed lines mark the contours of antennal discs, 2nd segments and 3rd segments, respectively. Arrowheads point to aristae. *peb-Gal4* is not expressed in the 3rd segment of larval antennal discs. At 0-4 h APF, *peb-Gal4* is expressed in a few cells in the 3rd segment. At 8-12 h APF, *peb-Gal4* expression is somewhat reduced. At 18 h APF and later, *peb-Gal4* is strongly expressed in all antennal cells. All images were acquired through confocal microscopy, except the one showing adult antennae, which was acquired from unfixed samples through fluorescence microscopy. Scale bars, 50 μm.

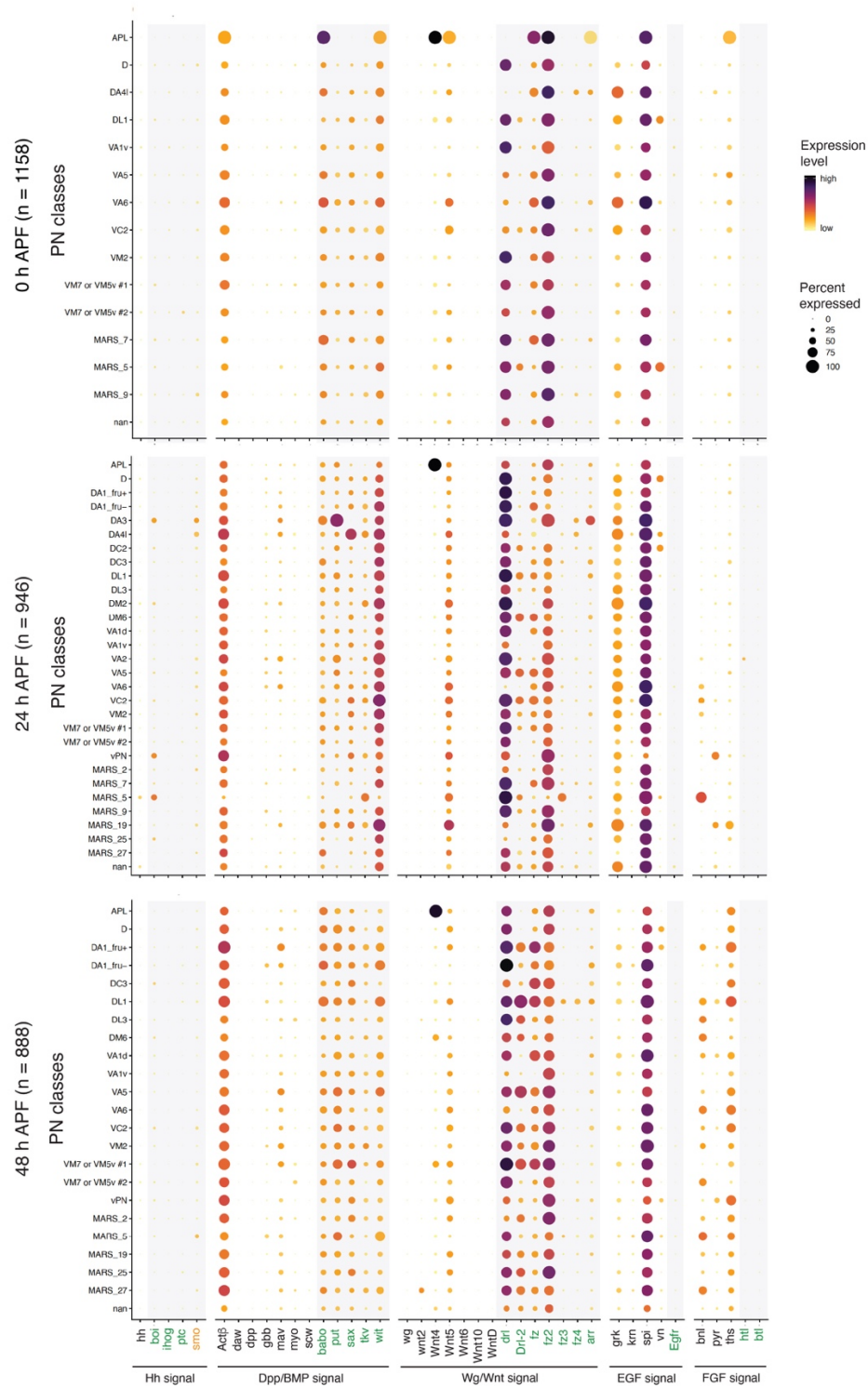

**Figure S4. Single-cell RNA-seq analyses of morphogen and morphogen receptor expression levels in PNs.**

Dot plots summarizing the expression of ligands and receptors in the Hh, TGFb, Wg/Wnt, EGF and FGF signaling pathways in PN types at 0 h APF, 24 h APF, 48 h APF and adult stage.

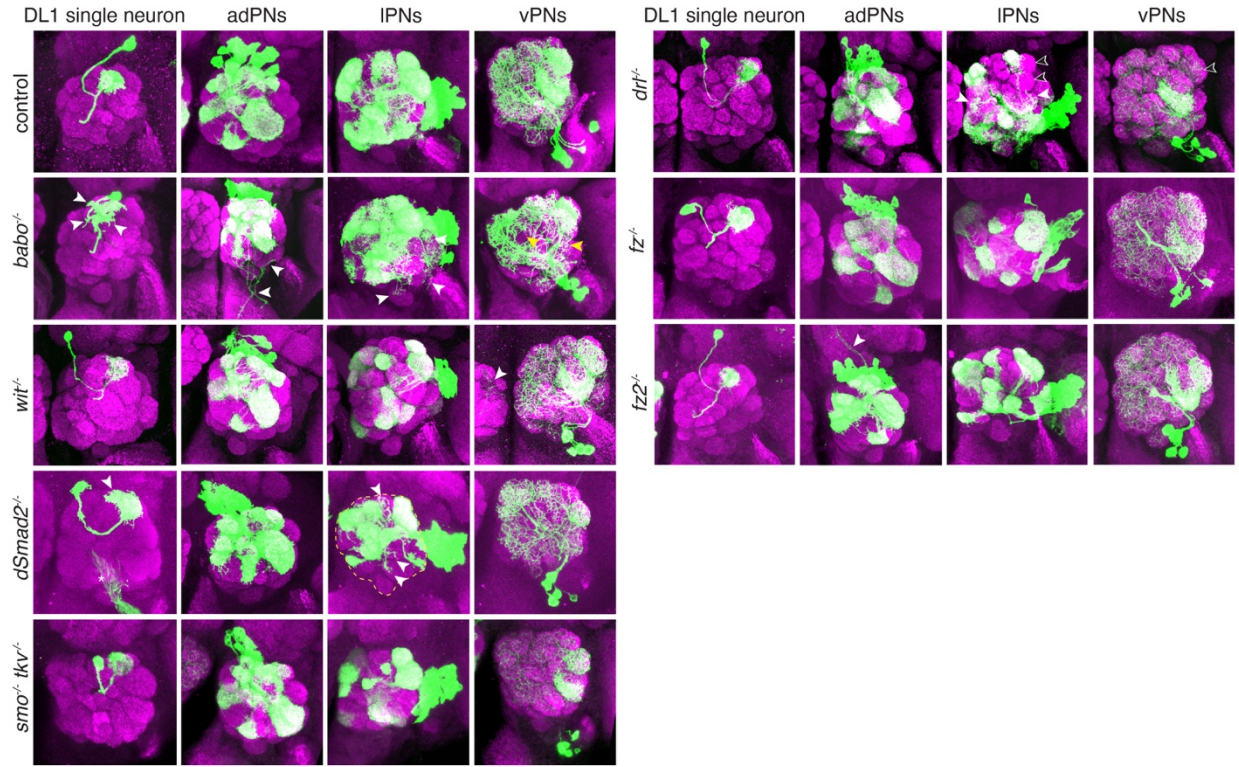

**Figure S5. Dendrite development of PNs carrying morphogen receptor and effector mutations.**

Representative confocal images of DL1 PN single cell clones, adPN neuroblast clones, IPN neuroblast clones and vPN neuroblast clones that carried different morphogen receptor mutations. White arrowheads indicate abnormal dendrite extensions.

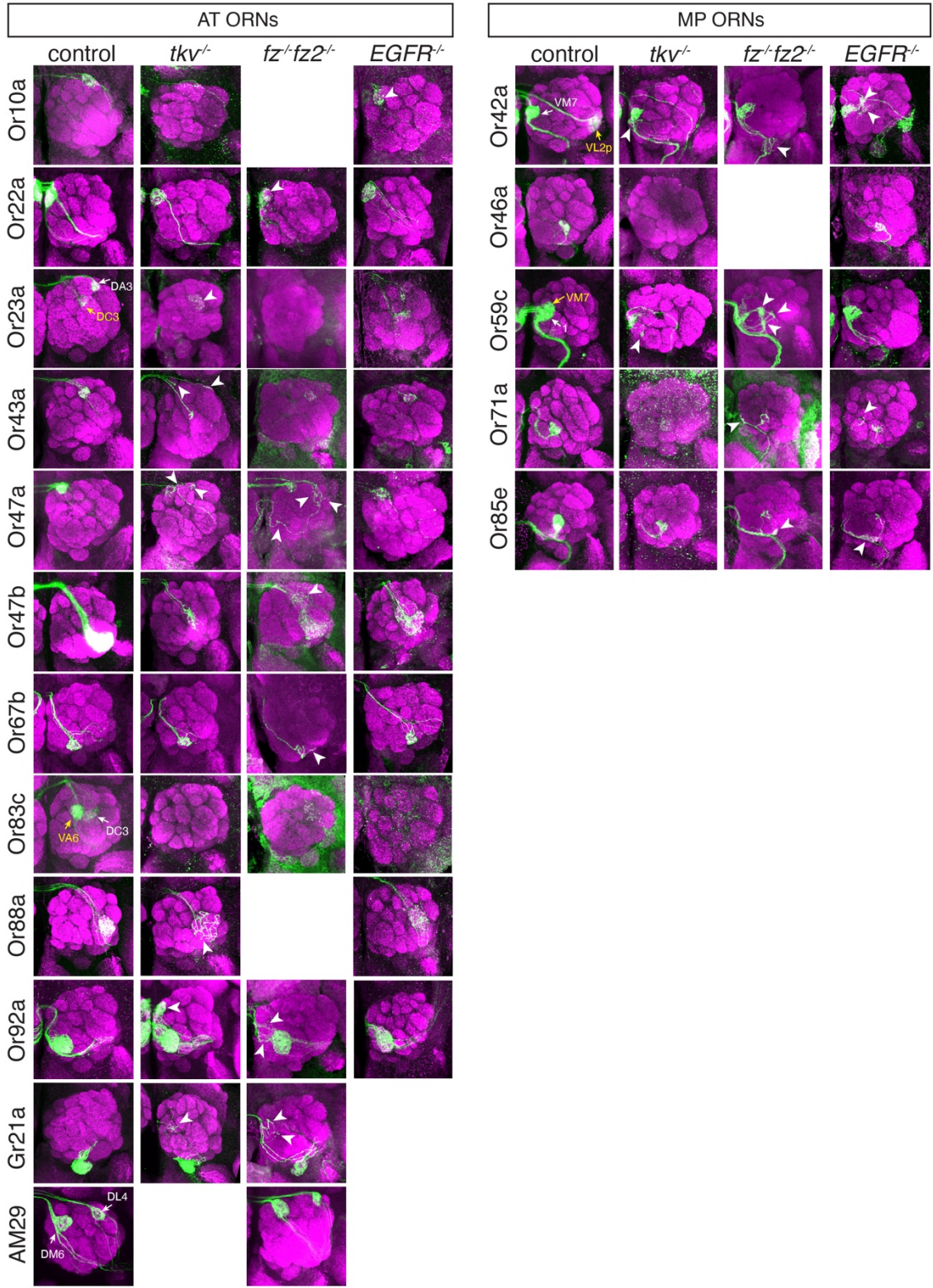

**Figure S6. Axonal targeting of ORNs carrying morphogen receptor and effector mutations.**
Representative confocal images showing axonal targeting of different ORN classes in the antennal lobes.
White arrowheads indicate mistargeted axons.

**Table S1. Genotypes of flies used in experiments described in Figures 1-6 and S1-S4.**

| **Figure** | **Genotype** |
| --- | --- |
| **Fig. 1B (3rd L)** | *w[\*] peb-GAL4; FRTG13 UAS-mCD8GFP* |
| **Fig. 1B (24 h APF)** | *(y[\*]) w[\*];ptc-GAL4/UAS-GFP-nls (14)/+;ry[506] p{PZ}hh[P30]/+* |
| **Fig. 1B (adult)** | *Canton-S* |
| **Fig. 1C** | *(y[\*]) w[\*];ptc-GAL4/UAS-GFP-nls (14)/+;ry[506] p{PZ}hh[P30]/+* |
| **Fig. 3 (*hh*, *dpp*, *wg*)** | *w[\*];UAS-mCD8GFP.1/+; ry[506] p{PZ}hh[P30]/p{GAL4-dpp.blk1}c40.1* |
| **Fig. 3 (*Activinβ-GFP*)** | *y[1] w[1118]; Mi{DH.1}Actbeta[MI14795-DH.PT-GFSTF.1]* |
| **Fig. 3 (*gbb-GAL4*)** | *w[\*]; P{GawB}NP2048 (gbb-GAL4)/ CyO* |
| **Fig. 3 (*bnl-GFP*)** | *y[1] w[\*]; Mi{PT-GFSTF.1}bnl[MI00874-GFSTF.1]/TM3, Sb[1] Ser[1]* |
| **Fig. 3 (*Wnt5-GAL4*)** | *y[\*] w[\*] P{GawB}NP6034 /+; UAS-mCD8GFP.1/CyO* |
| **Fig. 3 (*spi*)** | *(y[\*]) w[\*];ptc-GAL4/UAS-GFP-nls (14)/+;ry[506] p{PZ}bnl[06916]/+* |
| **Fig. 3 (*vn-GFP*)** | *y[1] w[\*]; Mi{PT-GFSTF.1}vn[MI05869-GFSTF.1]/TM6C Sb[1] Tb[1]* |
| **Fig. 4** | *w[\*] peb-GAL4; FRTG13 UAS-mCD8GFP* |
| **Fig. 5A (*hh*, *dpp*, *wg*)** | *w[\*];UAS-mCD8GFP.1/+; ry[506] p{PZ}hh[P30]/p{GAL4-dpp.blk1}c40.1* |
| **Fig. 5A (*Activinβ-GFP*)** | *y[1] w[1118]; Mi{DH.1}Actbeta[MI14795-DH.PT-GFSTF.1]* |
| **Fig. 5A (*gbb-GAL4*)** | *w[\*]; P{GawB}NP2048 (gbb-GAL4)/ CyO* |
| **Fig. 5A (*bnl-GFP*)** | *y[1] w[\*]; Mi{PT-GFSTF.1}bnl[MI00874-GFSTF.1]/TM3, Sb[1] Ser[1]* |
| **Fig. 5A (*Wnt5-GAL4*)** | *y[\*] w[\*] P{GawB}NP6034 /+; UAS-mCD8GFP.1/CyO* |
| **Fig. 5A (*vn-GFP*)** | *y[1] w[\*]; Mi{PT-GFSTF.1}vn[MI05869-GFSTF.1]/TM6C Sb[1] Tb[1]* |
| **Fig. 5C, fig. S3 (control)** | *hsFLP122/y w (or Y);FRTG13 tubp-GAL80/FRTG13 GH146-GAL4 UAS-mCD8GFP;;* |
| **Fig. 5C (*smo*)** | *y w hsFLP UAS-mCD8GFP/y w hsFLP (or Y); smo[3] FRT40A/tubp-GAL80 FRT40A GH146-GAL4;;* |
| **Fig. 5C (*ptc*)** | *y w hsFLP UAS-mCD8GFP/(y) w (or Y); FRT42D ptc[IIW] GH146-GAL4 UAS-mCD8GFP/FRT42D tubp-GAL80;;* |
| **Fig. 5C (*tkv*)** | *y w hsFLP UAS-mCD8GFP/y w hsFLP (or Y); tkv[*strII*] FRT40A/tubp-GAL80 FRT40A GH146-GAL4;;* |
| **Fig. 5C (*put*)** | *y w hsFLP UAS-mCD8GFP/y w hsFLP122 (or Y); GH146-GAL4 UAS-mCD8GFP/+;FRT82B tubp-GAL80/FRT82B put[135];* |
| **Fig. 5C (*EGFR*)** | *y w hsFLP UAS-mCD8GFP/(y) w (or Y); FRT42D EGFR[-] GH146-GAL4 UAS-mCD8GFP/FRT42D tubp-GAL80;;* |
| **Fig. 5C (*fz fz2*)** | *y w hsFLP122 UAS-mCD8GFP/y w hsFLP (or Y); GH146-GAL4 UAS-mCD8GFP/+;fz[H51] Dfz2[C1] FRT2A/ tubp-GAL80 FRT2A;* |
| **Fig. 6A** | (see genotypes shown for fig. S4 below) |
| **fig. S1** | *(y[\*]) w[\*];ptc-GAL4/UAS-GFP-nls (14)/+;ry[506] p{PZ}hh[P30]/+* |
| **fig. S3 (*babo*)** | *y w hsFLP UAS-mCD8GFP/y w (or Y); FRTG13 babo[Fd4] GH146-GAL4/FRTG13 tubp-GAL80;;* |
| **fig. S3 (*wit*)** | *y w hsFLP122 UAS-mCD8GFP/ w (or Y); GH146-GAL4 UAS-mCD8GFP/+; wit[G5] FRT2A/ tubp-GAL80FRT2A;* |
| **fig. S3 (*dSmad2*)** | *dSMAD2[-] UAS-mCD8GFP FRT19A/hsFLP122 tub-pGAL80 FRT19A; GH146GAL6 UAS-mCD8GFP/+;;* |
| **fig. S3 (*smo tkv*)** | *y w hsFLP UAS-mCD8GFP/y w (or Y); ); smo[2] tkv[*strII*] FRT40A/tubp-GAL80 FRT40A GH146-GAL4;;* |
| **fig. S3 (*drl*)** | *y w hsFLP UAS-mCD8GFP/y w (or Y); drl[343] FRT40A/tubp-GAL80 FRT40A GH146-GAL4 UAS-mCD8GFP/+;;* |
| **fig. S3 (fz)** | *y w hsFLP122 UAS-mCD8GFP/y w hsFLP (or Y); GH146-GAL4 UAS-mCD8GFP/CyO (or SP); fz[H5]1 FRT2A/ tubp-GAL80 FRT2A;* |
| **fig. S3 (fz2)** | *y w hsFLP122 UAS-mCD8GFP/y w hsFLP (or Y); GH146-GAL4 UAS-mCD8GFP/CyO (or SP); Dfz2[C1]1 FRT2A/ tubp-GAL80 FRT2A;* |
| **fig. S4 (control)** | *UAS-mCD8GFP eyFLP FRT19A/Or-GAL4; FRT40A/tub-GAL80 FRT40A; ; (OR-GAL4: Or83c-GAL4 (1), Or88a-GAL4 (X2db))* |
|  | *UAS-mCD8GFP eyFLP FRT19A/+; AM29-GAL4 FRT40A/tub-GAL80 FRT40A; ;* |
|  | *UAS-mCD8GFP eyFLP FRT19A/+; FRT40A/tub-GAL80 FRT40A; Or-GAL4/+ ; (OR-GAL4: Or10a-GAL4 (15), Or23a-GAL4, Or42a-GAL4, Or43a-GAL4 (18d), Or46a-GAL4, Or47a-GAL4, Or47b-GAL4 (15-6), Or59c-GAL4 (21/7), Or67b-GAL4 (B3960), Or85e-GAL4 (1.14.1), Or92a-GAL4 (38.1), Gr21a-GAL4 (D1))* |
|  | *UAS-mCD8GFP eyFLP FRT19A/+; FRT40A/tub-GAL80 FRT40A; Or-GAL4 UAS-mCd8GFP/+ (OR-GAL4: Or22a-GAL4, Or71a-GAL4)* |
| **fig. S4 (tkv)** | *UAS-mCD8GFP eyFLP FRT19A/Or-GAL4; tkv[*strII*] FRT40A/tub-GAL80 FRT40A; ; (OR-GAL4: Or83c-GAL4 (1), Or88a-GAL4 (X2db))* |
|  | *UAS-mCD8GFP eyFLP FRT19A/+; tkv[*strII*] FRT40A/tub-GAL80 FRT40A; Or-GAL4/+ ; (OR-GAL4: Or10a-GAL4 (14.1), Or23a-GAL4, Or42a-GAL4, Or43a-GAL4 (18d), Or46a-GAL4, Or47a-GAL4, Or47b-GAL4 (15-6), Or59c-GAL4 (21.7), Or67b-GAL4 (B3960), Or85e-GAL4 (1.14.1), Or92a-GAL4 (38.1), Gr21a-GAL4 (D1))* |
|  | *UAS-mCD8GFP eyFLP FRT19A/+; tkv[*strII*] FRT40A/tub-GAL80 FRT40A; Or-GAL4 UAS-mCD8GFP/+ (OR-GAL4: Or22a-GAL4, Or71a-GAL4)* |
| **fig. S4 (fz fz2)** | *UAS-mCD8GFP eyFLP FRT19A/Or59c-GAL4 (129t5.4); ; fz[H51] fz2[C1] FRT2A/tub-GAL80 FRT2A;* |
|  | *UAS-mCD8GFP eyFLP FRT19A/+; Or-GAL4/+; fz[H51] fz2[C1] FRT2A/tub-GAL80 FRT2A; (OR-GAL4: Or22a-GAL4 (14.2), Or23a-GAL4 (17.2), Or42a-GAL4 (48.3B), Or43a-GAL4 (27.5), Or47a-GAL4 (15.2), Or47b-GAL4 (15-8), Or67b-GAL4 (B3960.1), Or71a-GAL4 (30.3), Or83c-GAL4 (73.3B), Or85e-GAL4 (2.19.1), Or92a-GAL4 (62.4), Gr21a-GAL4 (9.323)(2), AM29-GAL4)* |
| **fig. S4 (EGFR)** | *UAS-mCD8GFP eyFLP FRT19A/Or-GAL4; FRT42D EGFR[-]/FRT42D tub-GAL80; ; (OR-GAL4: Or83c-GAL4 (1), Or88a-GAL4 (X2db))* |
|  | *UAS-mCD8GFP eyFLP FRT19A/+; FRT42D EGFR[-]/FRT42D tub-GAL80; Or-GAL4/+ ; (OR-GAL4: Or10a-GAL4 (15), Or22a (14.2), Or23a-GAL4, Or42a-GAL4 (43.4), Or43a-GAL4 (18d), Or46a-GAL4, Or47a-GAL4 (15.4), Or47b-GAL4 (15-6), Or59c-GAL4 (21.7), Or67b-GAL4 (B3960), Or85e-GAL4 (1.14.1), Or92a-GAL4 (38.1))* |
|  | *UAS-mCD8GFP eyFLP FRT19A/+; FRT42D EGFR[-]/FRT42D tub-GAL80; Or-71a-GAL4 UAS-mCD8GFP/+* |
